# Recurrent Emergence of an Antiviral Defense through Repeated Retrotransposition and Truncation of CHMP3

**DOI:** 10.1101/2021.04.27.441704

**Authors:** Lara Rheinemann, Diane Miller Downhour, Kristen A. Davenport, Alesia N. McKeown, Wesley I. Sundquist, Nels C. Elde

**Author notes:** Correspondence to (N.C.E.). These authors contributed equally (Lara Rheinemann, Diane Miller Downhour).

## Abstract

Most restriction factors recognize virus features to execute antiviral functions. In contrast, we discovered retroCHMP3, which instead impairs the host endosomal complexes required for transport (ESCRT) pathway to inhibit budding of enveloped viruses, including HIV-1. The ESCRT pathway is essential, so ESCRT inhibition creates the potential for cytotoxicity. We chart independent courses of retroCHMP3 emergence and reduction of cytotoxicity in New World monkeys and mice using ancestral reconstructions. Overexpression of full-length CHMP3 results in modest antiviral activity, which is enhanced by truncating mutations but causes increased cytotoxicity. We show that retroCHMP3 from squirrel monkeys acquired ancient mutations mitigating cytotoxicity before gaining the activating truncation. In contrast, a truncating mutation arose soon after the independent appearance of murine retroCHMP3, but the variant exhibits regulated expression by interferon signaling, illustrating distinct paths in the emergence of antiviral functions. RetroCHMP3 genes can repeatedly emerge in different species to independently create new immune functions.

## Introduction

Although many genes encoding immune defenses exhibit rapid evolution (Daugherty and Malik, 2012), most have ancient origins and are widely conserved among modern species. Mammalian hosts constantly adapt to infection with diverse viruses and evolved hundreds of immune genes that provide cell-autonomous protective responses, raising the possibility that some of these genes emerged recently in specific lineages or species.

One potential source of new genes is duplication of existing ones by retrotransposition of mRNA intermediates, a significant source of new genetic material (Kaessmann et al., 2009; Kubiak and Makalowska, 2017). This can occur when mobile genetic elements that generate copies of themselves by reverse transcription instead reverse transcribe cellular mRNAs and re-insert intronless retrocopies. In mammals, long interspersed element-1 (LINE-1) retrotransposons are the primary source of such retrocopies (Esnault et al., 2000; Gibbs et al., 2004; Lander et al., 2001; Mouse Genome Sequencing et al., 2002). Most retrocopies are “dead on arrival” as processed pseudogenes that degrade over time, but some retrocopies are expressed, often acquiring differential regulation and expression patterns (Rosso et al., 2008; Young et al., 2010; Yu et al., 2006), to become functional retrogenes. These retrogenes can replace the functions of parental genes (Ciomborowska et al., 2013; Kim et al., 2014) or diverge to encode novel functions (Bradley et al., 2004; Burki and Kaessmann, 2004; Young et al., 2010).

LINE-1-mediated retroduplication facilitated the emergence of several novel genes that function in antiviral immunity. In several primate species, independent retrotransposition of the coding sequence of cyclophilin A within the coding sequence of the restriction factor TRIM5α produced chimeric TRIMCyp genes that can potently block retrovirus replication (Brennan et al., 2008; Liao et al., 2007; Newman et al., 2008; Nisole et al., 2004; Sayah et al., 2004; Virgen et al., 2008; Wilson et al., 2008). Similarly, retroduplications of the retroviral restriction factors Fv1 in wild mice (Yap et al., 2020) and APOBEC3 in primates (Yang et al., 2020) appear to have expanded the capacity of these proteins to restrict retroviruses by increasing gene copy numbers and allowing functional divergence.

We recently discovered an inhibitor of enveloped virus budding, retroCHMP3, which arose independently in house mice and New World monkeys by LINE-1-mediated retrotransposition and subsequent C-terminal truncation of the ESCRT-III protein CHMP3 (Rheinemann et al., 2020). The ESCRT pathway mediates essential cellular membrane remodeling events, including multivesicular body formation, cytokinetic abscission, and resealing of the post-mitotic nuclear envelope (Christ et al., 2017; Henne et al., 2013; McCullough et al., 2018; Scourfield and Martin-Serrano, 2017). Many enveloped viruses hijack the ESCRT-pathway to bud from cellular membranes (Votteler and Sundquist, 2013) and some quasi-enveloped viruses also depend on the ESCRT pathway for non-lytic release (Feng et al., 2014; Himmelsbach et al., 2018; McKnight and Lemon, 2018). Since the ESCRT pathway is exploited by many virus families, inhibition of ESCRT functions during budding has the possibility to block the replication of diverse viruses. However, because the ESCRT pathway functions during essential cellular events such as cell division, loss of ESCRT function is typically cytotoxic.

Sensitivity of the ESCRT pathway to perturbation results from complex interactions between ESCRT-III proteins such as CHMP3, which form membrane-bound filaments of interlocked subunits. Following filament formation, the AAA ATPase VPS4 dynamically extracts individual ESCRT-III subunits to remodel the filaments, which progressively draws the bound membranes together and facilitates membrane fission (Caillat et al., 2019; McCullough et al., 2018; Remec Pavlin and Hurley, 2020). Like all ESCRT-III proteins, the N-terminal end of CHMP3 contains the core polymer-forming unit, while the C-terminal end contains an autoinhibitory domain and binding sites for VPS4 (McCullough et al., 2018). Therefore, engineered C-terminally truncated versions of CHMP3 that lack these functional elements act as strong dominant-negative inhibitors of ESCRT functions, including virus budding and cytokinesis (Dukes et al., 2008; Zamborlini et al., 2006). In contrast, naturally occurring retroCHMP3 proteins broadly inhibit the budding of diverse enveloped virus families by impairing ESCRT-mediated membrane fission (**Fig. 1A**) but exhibit remarkably little toxicity and do not block cellular ESCRT functions during abscission. Hence, retroCHMP3 proteins have evolved to spare cellular ESCRT functions while potently inhibiting ESCRT-dependent virus budding (Rheinemann et al., 2020).

**Fig. 1:**
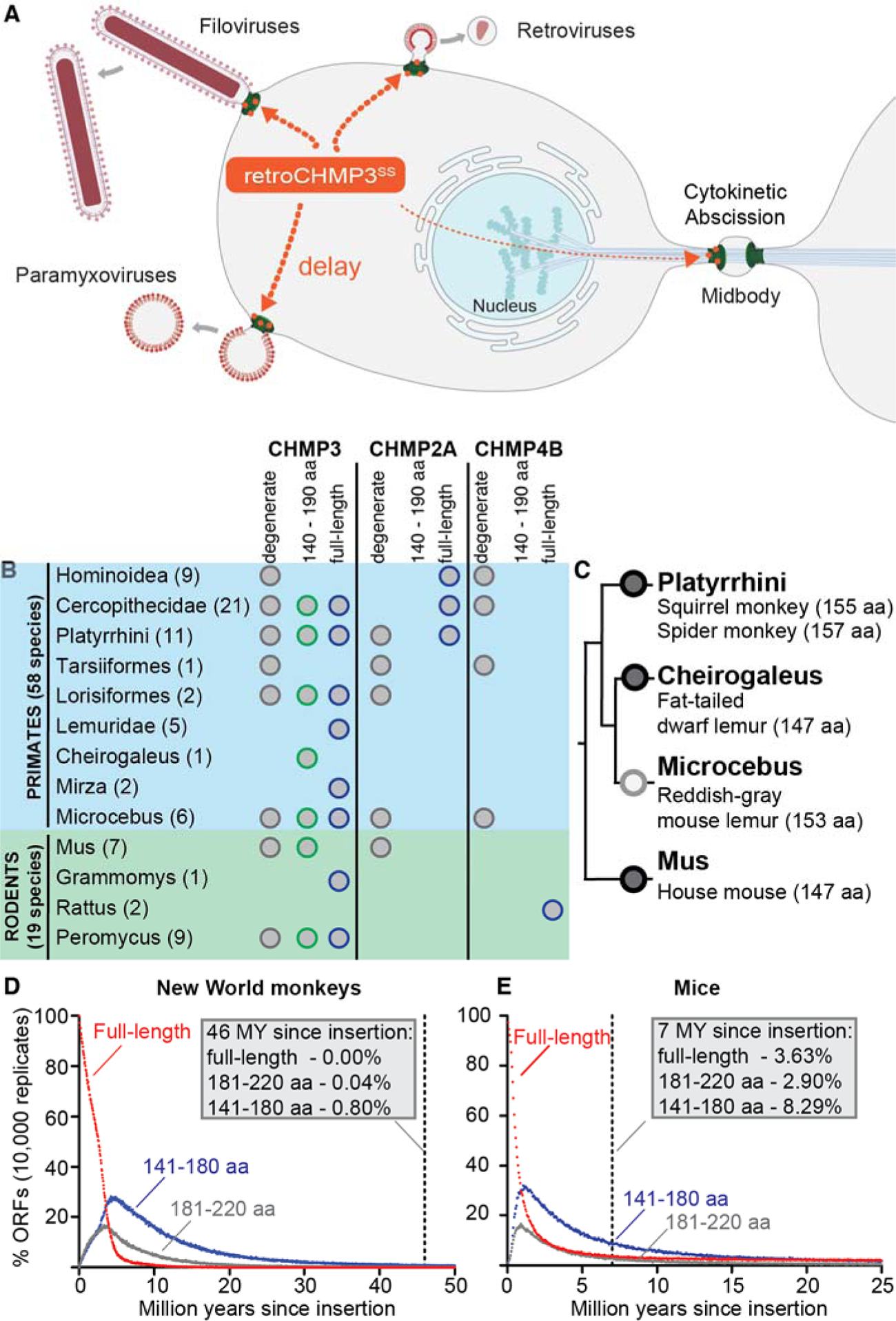
CHMP3 retrocopies are frequently observed in primate and rodent genomes. (A) RetroCHMP3 proteins potently inhibit budding of ESCRT-dependent viruses while only modestly affecting essential ESCRT-dependent cellular processes like cytokinetic abscission. **(B)** CHMP3 retroduplication is common in primates and rodents. Species were analyzed for the presence of at least one retrocopy of CHMP3, CHMP2A, and CHMP4B by NCBI BLAST of whole-genome shotgun contigs, and data were collapsed into families. Numbers behind each family name indicate the number of available genomes per family. For a complete list of species investigated and details of identified CHMP3 retrocopies, see Tables S1 and S2. **(C)** Independent acquisition of truncated retrocopies of CHMP3 of ∼150 amino acids length. Circles denote retrotransposition of full-length CHMP3 in common ancestors of New World monkeys (Platyrrhini), dwarf lemurs (Cheirogaleus), mouse lemurs (Microcebus), and house mice (Mus). Species names indicate species that acquired premature stop codons, leading to C-terminally truncated predicted protein products of ∼150 amino acids length. Dark grey circles denote retrocopies confirmed by sequencing of genomic DNA. Light grey circle denotes retrocopy detected by NCBI BLAST of whole-genome shotgun contigs. (**D**), (**E**) Simulation of open reading frame (ORF) loss of CHMP3 retrocopies using mutation rates and generation times for New World monkeys **(D)** and mice **(E)** given a model of relaxed constraint (neutral evolution). Dots represent the percentage of simulated ORFs still intact after a given time. Vertical dashed lines indicate estimated current age of the two retroCHMP3 genes.

Here we report evolutionary courses of retroCHMP3 emergence and reduction of cytotoxicity in primates and rodents. We discovered repeated, independent duplications of CHMP3 by LINE-1 mediated retrotransposition in multiple species. The resulting CHMP3 retrocopies persist more frequently than expected over millions of years of evolution. In several primate species, the full-length CHMP3 open reading frame (ORF) was retained, and full-length CHMP3 proteins predicted to be encoded by these retrogenes became progressively less cytotoxic but retained the ability to inhibit virus budding. When artificially truncated by removing roughly 70 C-terminal amino acids, these retrocopies became potent virus budding inhibitors, suggesting they are only a single truncating mutation away from being powerful antiviral proteins. In mice, retroCHMP3 expression is modestly interferon-inducible, suggesting regulation on the transcriptional level. The independent retroduplication and neofunctionalization of CHMP3 in multiple mammalian species reveal a potentially versatile means for acquiring antiviral functions by restricting core cellular functions exploited by viruses.

## Results

### CHMP3 retrocopies frequently persist in primate and rodent genomes

At least three mammalian species independently acquired LINE-1-mediated duplications of CHMP3 (retroCHMP3) that inhibit virus budding, including house mice and two species of New World monkeys (Rheinemann et al., 2020). To determine the frequency of retroCHMP3 protein acquisition compared to other CHMP paralogs, we searched for CHMP3 retrocopies in a larger sample of available primate and rodent genomes. CHMP3 retrocopies were identified by manual BLAST search (Altschul et al., 1990) of the National Center for Biotechnology Information (NCBI) database of whole-genome shotgun contigs. Specifically, we searched for DNA sequences with high similarity to the human (for primate genomes) or mouse (for rodent genomes) CHMP3 mature RNA transcript and lacking introns, which is a hallmark of LINE-1-mediated retrotransposition. This search identified several primate and rodent genomes containing one or more CHMP3 retrocopies (**Fig. 1B, Table S1, S2**). Based on the surrounding genomic context and differences in flanking target-site duplications, we determined that the CHMP3 retrocopies in different species were generated by a number of independent evolutionary events (**Table S3**).

Many CHMP3 retrocopies acquired truncating stop codons that result in predicted ORFs of 140 to 190 amino acids (**Fig. 1B**, green circles). Such retrocopies encode proteins that are similar in length to the well-characterized retroCHMP3 proteins in squirrel monkeys and mice (Rheinemann et al., 2020). These retrocopies are expected to be potent inhibitors of ESCRT function due to the deletion of VPS4 binding sites and autoinhibitory sequences (Bajorek et al., 2009; Lata et al., 2008; Muziol et al., 2006; Shim et al., 2007; Stuchell-Brereton et al., 2007; Zamborlini et al., 2006), and therefore have the potential to function as virus budding inhibitors. Five out of nine primate families and two out of four rodent families contained truncated retrocopies of 140–190 amino acids, in most cases in multiple species per family (**Fig. 1B, Table S3**). In addition to three previously reported retroCHMP3 proteins in New World monkeys and mice, which are truncated to approximately 150 amino acids (Rheinemann et al., 2020), we identified two CHMP3 retrocopies with similar truncation lengths in mouse lemurs and dwarf lemurs (**Fig. 1C**). In contrast to the high prevalence of truncated retroCHMP3 retrocopies, we did not find any truncated retrocopies of the two paralogous ESCRT-III proteins CHMP2A and CHMP4B in the same primate and rodent genomes (**Fig. 1B**).

We also identified a large number of full-length retrocopies of CHMP3 (**Fig. 1B**, blue circles). These genes resulted from recent duplication events or from long-term persistence of retrogenes in which any acquired mutations do not disrupt the original ORF. Six out of nine primate families and two out of four rodent families contained full-length CHMP3 retrocopies. As expected, we also found many degenerate CHMP3 retrocopies (**Fig. 1B**, grey circles) which had acquired multiple truncating nonsense mutations. Full-length retrocopies of CHMP2A and CHMP4B, in contrast, were rare, and most CHMP2A and CHMP4B retrocopies were highly degraded. These observations indicate that CHMP3 retrocopies are either more frequently generated than CHMP2A and CHMP4B retrocopies and/or persist much longer as full-length and truncated retrocopies.

To determine whether retroCHMP3 retrocopies persist at a higher rate than predicted by chance, we modeled the anticipated temporal degradation of full-length CHMP3 retrocopies in the absence of selection pressure using estimates for mutation rates and generation times for New World monkeys **(Fig. 1D**) and mice (**Fig. 1E**). The probability of retaining a full-length CHMP3 retrocopy under relaxed constraint fell below 5% within 5.51 MY and 4.55 MY in New World monkeys and mice, respectively. We estimate that the retroCHMP3 copy in New World monkeys was inserted 43-46 MYA based on the syntenic copy found in all New World monkey species examined, which is absent in hominoids and Old World monkeys (**Fig. S1A**) (Chatterjee et al., 2009; Finstermeier et al., 2013; Perelman et al., 2011; Pozzi et al., 2014). The probability of survival of a full-length, or nearly full-length, CHMP3 retrocopy surviving after 46 MY was 0.00% and 0.04%, respectively. The probability of persistence of a CHMP3 retrocopy of 141-180 amino acids length was 0.80%.

Following similar analyses, we estimate that the retroCHMP3 copy in mice was inserted 7 MYA based on a syntenic copy found in several Mus genomes, which is absent in *Mus pahari* (**Fig. S1B**) (Steppan and Schenk, 2017). In our simulations, only 3.63% of retrocopies did not acquire a premature stop codon after 7 MY, and 2.90% and 8.29% retained an ORF of at least 181 and 141 amino acids, respectively. Therefore, the odds of CHMP3 retrocopies persisting in full-length or truncated forms in the absence of positive selection over the long periods is very low in primates and low in rodents. Together with the paucity of viable retrocopies of CHMP2 and CHMP4, these observations indicate that retroCHMP3 proteins confer an evolutionary advantage under intermittent or continuous selection in primates and mice.

### Overexpression of full-length CHMP3 retrocopies moderately inhibits virus budding

To characterize the antiviral activities of different types of CHMP3 retrocopies, we focused on the single origin CHMP3 retrocopy found in New World monkeys. This CHMP3 retrocopy arose in a common ancestor of New World monkeys but underwent various fates in different species (**Fig. 2A, S2A, Table S4**): For two species in our sample, squirrel monkeys and spider monkeys, CHMP3 retrocopies were independently truncated to encode proteins of 155 amino acids (squirrel monkeys) and 157 amino acids (spider monkeys), and the resulting proteins can both potently inhibit budding of ESCRT-dependent enveloped viruses (Rheinemann et al., 2020). In titi, saki, capuchin, and red howler monkeys, the CHMP3 retrocopies acquired pseudogenizing mutations that severely disrupt the ORF. In contrast, owl monkeys and pygmy marmosets encode full-length CHMP3 ORFs, while woolly monkeys gained 24 amino acids at the carboxy terminus of CHMP3 owing to loss of the original stop codon (**Fig. 2A**). Therefore, New World monkey retrocopies of CHMP3 make it possible to study and compare the impact of different evolutionary fates on the function of retroCHMP3 proteins.

**Fig. 2:**
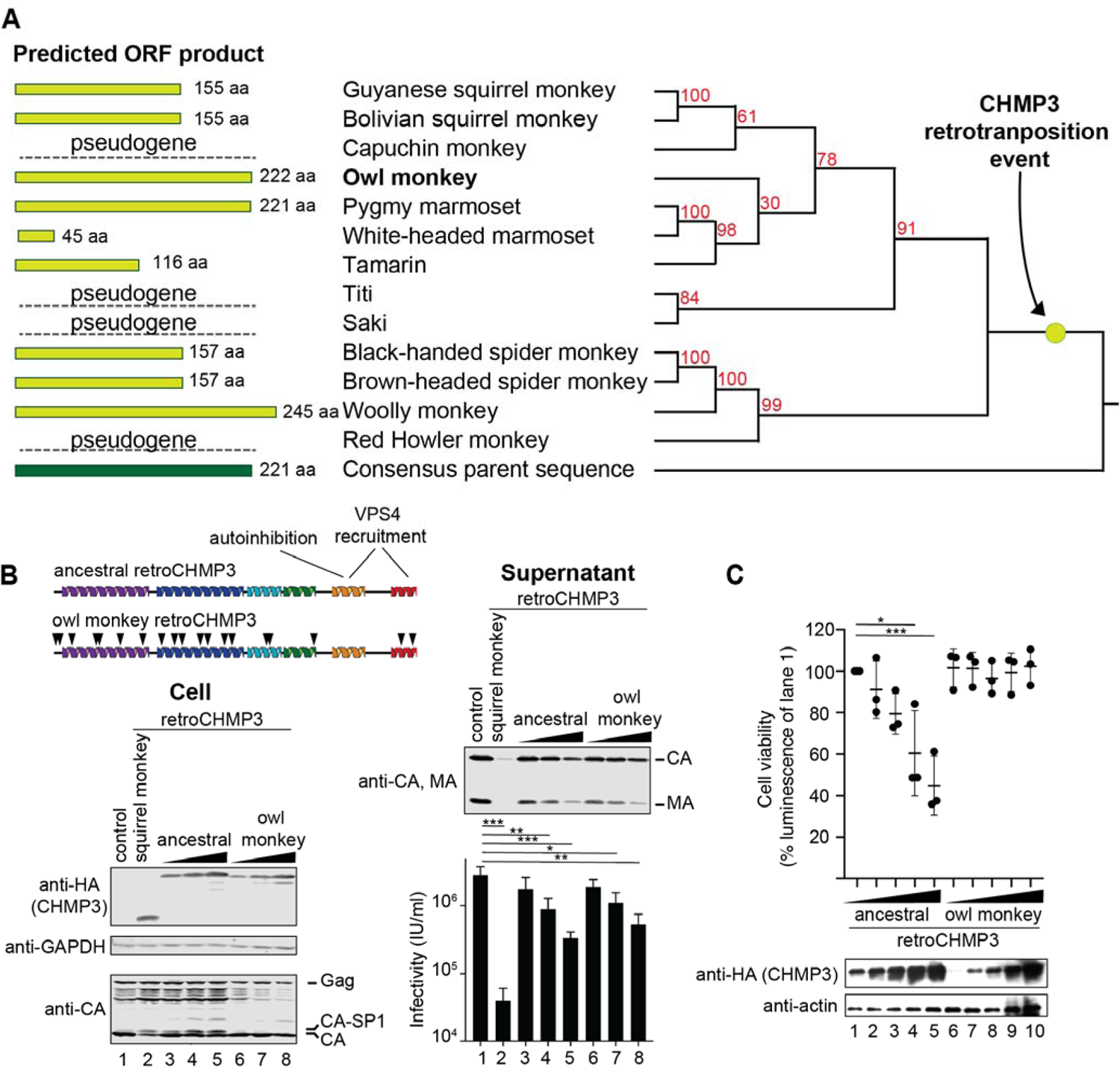
Overexpression of full-length CHMP3 retrocopies moderately inhibits virus budding. **(A)** Phylogeny of retroCHMP3 in New World monkeys. Light green bars show the length of predicted protein products of modern retroCHMP3 ORFs. The highest confidence consensus sequence of parental CHMP3 genes (consensus parent) was used as an approximation of the ancestral CHMP3 retrocopy and depicted as the outgroup. Red numbers indicate bootstrap support for each node. See also Fig. S2. Nucleotide and amino acid sequences of retroCHMP3 in modern New World monkeys are given in Table S4. **(B)** Transient overexpression of HA-tagged ancestral retroCHMP3 and owl monkey retroCHMP3 in human embryonic kidney (HEK293T) cells moderately inhibits the release of HIV-1 proteins (MA, CA) and reduces viral titers in the supernatant. Titer graphs show mean ± SD from 3 experimental replicates Schematic shows ancestral retroCHMP3 and modern owl monkey retroCHMP3. Black arrowheads indicate amino acid substitutions relative to ancestral retroCHMP3. **(C)** Viability of cells transiently transfected with HA-tagged ancestral retroCHMP3 and owl monkey retroCHMP3, as determined in a luminescent ATP cell viability assay. Mean ± SD from ≥ 3 experimental replicates with n 6 each. Here and throughout, **** p<0.0001, *** p<0.001, **p<0.01, *p<0.05 by one-way ANOVA followed by Tukey’s multiple comparisons test. For clarity, only comparisons with lane 1 are shown. For a complete set of pairwise comparisons in this and all following figures, see Table S6.

While truncated variants of retroCHMP3 inhibit ESCRT-dependent virus budding, it isn’t known whether full-length CHMP3 can do the same. We hypothesized that the ancestral full-length CHMP3 retrocopy might have provided at least modest antiviral activity to explain the otherwise improbable retention of full-length or truncated CHMP3 retrocopies over 46 MY. To test this idea, we reconstructed the ancestral retrocopy using nucleotide sequences of modern parental CHMP3 and CHMP3 retrocopies in New World monkeys (**Fig. 3A, Table S4)**. A high level of conservation for the parental CHMP3 gene among primates facilitated the generation of a high-quality inferred ancestral sequence. The full-length ancestral CHMP3 retrocopy inhibited HIV-1 budding modestly but significantly when transiently expressed in human embryonic kidney (HEK293T) cells (**Fig. 2B**). Intriguingly, the modern full-length retroCHMP3 protein from owl monkeys also inhibited HIV-1 budding to a similar extent as the ancestral retrocopy, despite having acquired 19 amino acid substitutions in comparison to the ancestral CHMP3 retrocopy (**Fig. 2B, S3**). These results demonstrate that full-length CHMP3 proteins can inhibit virus budding, perhaps by altering the optimal stoichiometry of subunits in ESCRT-III filaments. This observation raises the possibility that a moderate inhibitory effect on virus budding conferred a selective advantage to hosts and disfavored pseudogenization even over long evolutionary intervals.

**Fig. 3:**
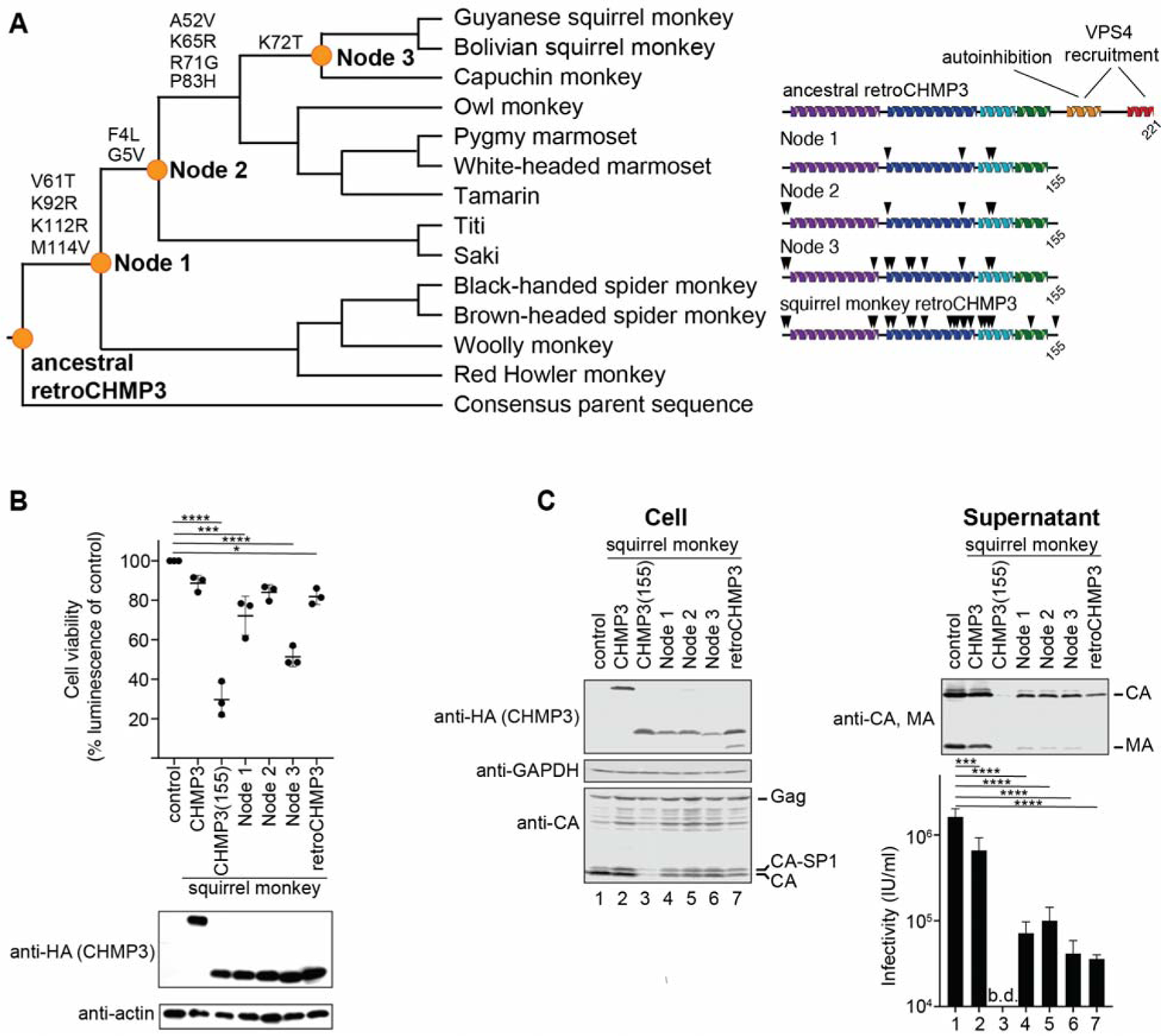
Ancestral intermediates of squirrel monkey retroCHMP3 exhibit gains and losses of cytotoxicity. (A) Reconstruction of ancestral retroCHMP3 and intermediates of squirrel monkey retroCHMP3 (orange dots). Amino acid substitutions leading to modern squirrel monkey retroCHMP3 are listed over branches. Schematic shows ancestral retroCHMP3, artificially truncated ancestral intermediates, and squirrel monkey retroCHMP3. Black arrowheads indicate amino acid substitutions relative to the ancestral retroCHMP3. Nucleotide and amino acid sequences of ancestral intermediates are shown in Table S4. **(B)** Viability of cells transiently transfected with indicated HA-tagged constructs, as determined in a luminescent ATP cell viability assay. Mean ± SD from 3 experimental replicates with n ≥ 6 each. **(C)** Transient overexpression of HA-tagged, prematurely truncated ancestral intermediates in human embryonic kidney (HEK293T) cells inhibits the release of HIV-1 proteins (MA, CA) and reduces viral titers in the supernatant. Titer graphs show mean ± SD from 3 experimental replicates. b.d. – below detection limit.

When highly overexpressed, the ancestral retroCHMP3 protein is also mildly cytotoxic (**Fig. 2C**), indicating that full-length retroCHMP3 proteins can also exact a cost to cell viability. In contrast, similar high levels of owl monkey full-length retroCHMP3 protein did not induce cytotoxicity. This remarkable observation suggests that the modest cytotoxicity of the ancestral CHMP3 retrocopy may have favored the accumulation of mutations that reduced this cytotoxicity while retaining modest antiviral function in its full-length form. As a result, modern owl monkey retroCHMP3 maintained the capacity to inhibit virus budding while evolving to lose cytotoxicity.

### Ancestral intermediates of retroCHMP3 exhibit both gains and losses of cytotoxicity

RetroCHMP3 proteins appear to balance the selective advantage of virus inhibition with the cost of interfering with essential cellular ESCRT functions by acquiring mutations that reduce cytotoxicity but not inhibition of virus budding. To trace the reduction of cytotoxicity of ancestral retroCHMP3 over time, we reconstructed ancestral intermediates along the lineage to modern squirrel monkey retroCHMP3 protein (**Fig. 3A, S3**). After diverging from closely related capuchin monkeys, retroCHMP3 of squirrel monkeys acquired a truncating stop codon at 155 amino acids, which converts the protein into a potent inhibitor of virus budding (Rheinemann et al., 2020). Remarkably, the squirrel monkey protein is significantly less cytotoxic when compared to C-terminally truncated parental CHMP3 (**Fig. 3B**). To chart the evolutionary course of loss of cytotoxicity, we generated three ancestral sequences (**Fig. 3A**, nodes 1-3, orange dots) and truncated them at 155 amino acids to experimentally model the acquisition of an ancient activating mutation (**Fig. 3B**). Mutations acquired between the retroduplication event and nodes 1 and 2 led to sharp reductions in cytotoxicity, with node 2 comparable to modern squirrel monkey retroCHMP3. Truncated proteins corresponding to both nodes 1 and 2 potently inhibit HIV-1 budding (**Fig. 3C**), indicating that the amino acid substitutions did not impair the ability to inhibit virus budding. RetroCHMP3 from species descending from node 2 underwent a variety of different fates (**Fig. 2A, 3A**), from pseudogenization (titi, saki, and capuchin monkey), to retention of a full-length copy (owl monkey and pygmy marmoset), and premature truncation at various lengths (white-headed marmoset, tamarin, and squirrel monkey). In contrast, changes acquired between node 2 and node 3 increased the cytotoxicity of node 3 compared with node 2 (**Fig. 3B**) while still retaining efficient HIV-1 budding inhibition (**Fig. 3C**). Descendants of node 3 were either pseudogenized (capuchin) or accumulated additional mutations that further reduced cytotoxicity (squirrel monkey). These results are consistent with randomly occurring mutations followed by selection to retain variants with at least modest antiviral activity and reduced cytotoxicity. CHMP3 appears to diversify with many potential substitutions that mitigate cytotoxicity. Of particular note, it appears the first mutations (Node 1) acquired in retroCHMP3 from ancient New World monkeys drastically reduced cytotoxicity of the truncated variant, which set the stage for the dynamic history of retroCHMP3 activation and loss observed in modern species.

### Modern full-length retroCHMP3 proteins are poised to become potent virus budding inhibitors

Prior to acquiring truncating mutations, full-length retroCHMP3 proteins only modestly inhibit virus budding. In our sample of New World monkeys, squirrel monkeys and spider monkeys independently acquired premature stop codons that activate retroCHMP3 proteins into more potent inhibitors of virus budding (**Fig. 4A** and (Rheinemann et al., 2020)). To determine whether full-length retroCHMP3 proteins from other species can similarly be converted into potent virus budding inhibitors, we introduced truncating stop codons into the ORFs of retroCHMP3 encoded by woolly monkeys, owl monkeys, and pygmy marmosets. In all three cases, the engineered truncations of retroCHMP3 strongly inhibited HIV-1 budding, with potencies that were comparable to retroCHMP3 from squirrel monkeys (**Fig. 4A**). Importantly, these truncated retroCHMP3 proteins from woolly monkey, owl monkey, and pygmy marmoset caused little to no detectable cytotoxicity compared to a similarly truncated CHMP3 protein (CHMP3(155); **Fig. 4B**). These results reveal that several species of New World monkeys are one mutation away from encoding retroCHMP3 variants with reduced cytotoxicity that potently inhibit the budding of ESCRT-dependent viruses like HIV-1.

**Fig. 4:**
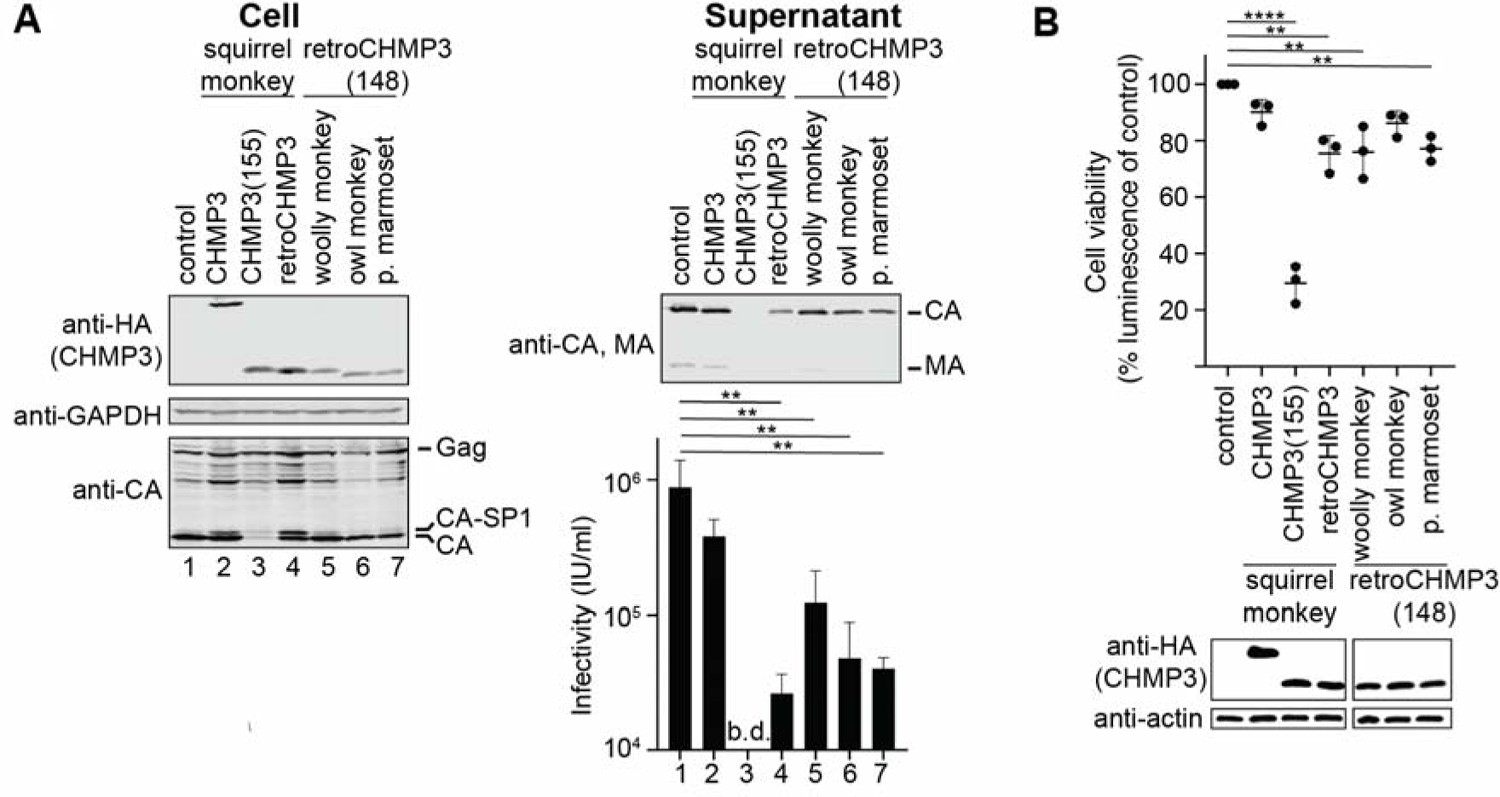
Modern full-length retroCHMP3 proteins are poised to become potent virus budding inhibitors. **(A)** Transient overexpression of truncated and HA-tagged retroCHMP3 variants from woolly monkeys, owl monkeys, and pygmy marmosets in human embryonic kidney (HEK293T) cells inhibits the release of HIV-1 proteins (MA, CA) and reduces viral titers in the supernatant. Titer graphs show mean ± SD from 3 experimental replicates. b.d. – below detection limit. **(B)** Viability of cells transiently transfected with indicated HA-tagged constructs, as determined in a luminescent ATP cell viability assay. Mean ± SD from 3 experimental replicates with n ≥ 6 each.

### Toxicity of the less ancient mouse retroCHMP3 toxicity may be controlled by interferon stimulation

Next we turned to the emergence of retroCHMP3 in mice, an independently acquired variant of retroCHMP3 that can also potently inhibit virus budding and is induced modestly by interferon signaling (Rheinemann et al., 2020). The murine CHMP3 retrocopy arose by LINE-1-mediated duplication into an intergenic region of *Mus musculus* chromosome 18, which contains several repetitive genetic elements, including LINEs, SINEs, and mouse endogenous retroviruses (MERVs; **Fig. S1B**). We confirmed the presence of a syntenic CHMP3 retrocopy in several *Mus* species by sequencing of genomic DNA (**Table S5, Fig. S1B**). For species where genomic DNA was not available, we examined the region in *Mus* genomes available on the UCSC Genome browser. Based on estimates of divergence times between ancestral species of mice and the absence of murine retroCHMP3 in *Mus pahari*, this retrocopy likely arose between 6–8 MYA (**Fig. 5**). A truncating stop codon resulting from a frameshift mutation and resulting in an ORF encoding 147 amino acids appeared relatively soon after the emergence of murine retroCHMP3, based on its absence in *Mus minutoides*. Therefore, in contrast to retroCHMP3 proteins in New World monkeys, which became less cytotoxic before acquiring activating stop codons, retroCHMP3 proteins in mice had substantially less time to sample mutations, even when accounting for the shorter generation times of mice compared to primates. Consistent with the relatively early acquisition of an activating stop codon, we previously found that mouse retroCHMP3 remains significantly more cytotoxic than squirrel monkey retroCHMP3 and is induced modestly by interferon signaling, presumably by putative interferon-responsive elements in the upstream region of murine retroCHMP3 (Rheinemann et al., 2020). Two putative interferon-responsive STAT1/STAT3 binding sites are present in *Mus pahari* and therefore predate the emergence of retroCHMP3 (**Fig. 5**). In addition, a murine retroviral related sequence (MuRRS) element is found 5.5 kB upstream of the retroCHMP3 ORF in species closely related to *Mus musculus*, which also may have conferred expression regulation through interferon-induced signaling (Chuong et al., 2017).

**Fig. 5:**
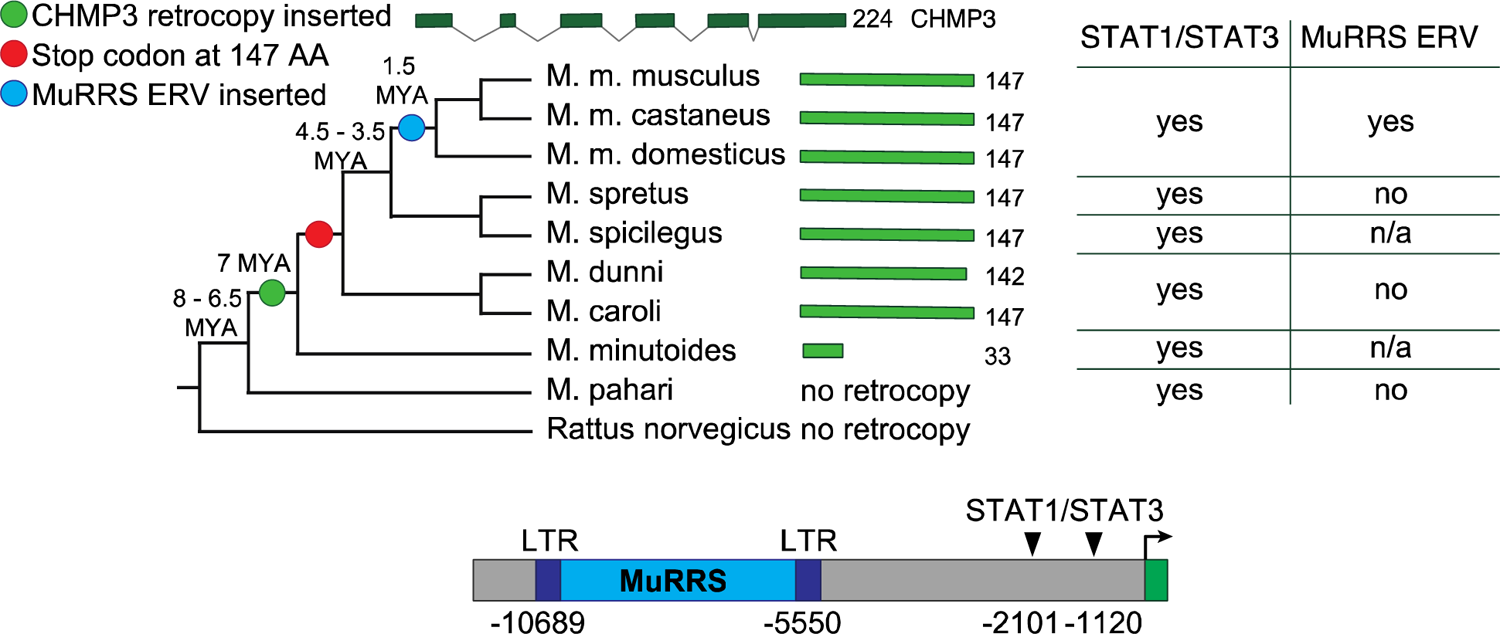
Expression of mouse retroCHMP3 can be controlled by interferon signaling. Phylogeny of Mus genera is based on (Steppan and Schenk, 2017). Light green bars show length of modern retroCHMP3 copies. Table indicates presence of putative regulatory elements in Mus species. n/a – genome assembly not of sufficient quality to determine presence of MuRRS. Nucleotide and amino acid sequences and of retroCHMP3 in modern Mus species are listed in Table S5. Schematic shows putative regulatory elements upstream of retroCHMP3 ORF in Mus musculus. Numbers indicate nucleotides relative to retroCHMP3 start codon. LTR – long terminal repeat, MuRRS – murine retrovirus-related sequence, ERV – endogenous retrovirus.

Thus, in contrast to retroCHMP3 in New World monkeys, which acquired mutations that reduced cytotoxicity before gaining an activating mutation, a relatively cytotoxic truncated variant in mice seems to have persisted because low basal expression levels and modest interferon induction minimized cytotoxicity. Analysis of mouse and New World monkey retroCHMP3 variants therefore revealed two pathways to generate antiviral functions against diverse ESCRT-dependent viruses.

## Discussion

Lineage- and species-specific variants of retroCHMP3 are notable additions to cell-autonomous immune responses that interfere with different steps in virus replication cycles (Chemudupati et al., 2019; Doyle et al., 2015; Duggal and Emerman, 2012; Kluge et al., 2015). Many genes encoding antiviral proteins evolve rapidly in recurrent genetic conflicts with viruses (Daugherty and Malik, 2012) and also tend to be conserved across vertebrates, indicating ancient origins as immune defenses. In contrast, retroCHMP3 illustrates how immune responses might be regularly augmented with new defenses in different species.

In sampling primates and rodents, we discovered several independent origins of retroCHMP3 copies (**Fig. 1B**). Importantly, these copies persist as full-length or truncated variants far more frequently than predicted by chance (**Fig. 1D, E**) or than was observed for two functionally associated ESCRT-III paralogs, CHMP2 and CHMP4 (**Fig. 1b**). These observations provide strong evidence that CHMP3 retrocopies is positively selected for antiviral activity against ESCRT-dependent enveloped viruses (Rheinemann et al., 2020). Due to the broad reliance of virus families on the ESCRT pathway, CHMP3 retrogenes could provide broad inhibitory activity against the budding of diverse virus classes.

In addition, turnover of retroCHMP3 revealed by its patchy distribution among mammalian species implies that there are also substantial costs to altering ESCRT activity. We show that the ancestral full-length CHMP3 retrocopy in New World monkeys is indeed modestly cytotoxic (**Fig. 2**), which initially may have dictated low-level expression, consistent with observed low baseline expression of modern retroCHMP3 in squirrel monkey and mouse cell lines (Rheinemann et al., 2020). However, cytotoxicity can be drastically reduced by multiple different mutations in full-length or truncated CHMP3 retrocopies (**Fig. 2, 3**), and reduction of cytotoxicity can be achieved without losing virus budding inhibition (**Fig. 4**). Additionally, cytotoxicity may have been mitigated by transcriptional control, *e.g.,* modest interferon stimulation in mice (**Fig. 5**, (Rheinemann et al., 2020)). These observations suggest that: 1) CHMP3 retrocopies impose fitness costs on the host that can drive reduction of cytotoxicity, 2) in some species, selective pressure by viruses favored amino acid substitutions that balanced reduction of cytotoxicity with virus inhibition over pseudogenization, and 3) in the absence of such selective pressures, CHMP3 retrocopies eventually turn over either by genetic drift or purifying selection to alleviate cytotoxicity.

RetroCHMP3 proteins are among a handful of immune defenses that originate as rare byproducts of LINE-1 retrotransposition. In some wild mouse species, retroduplication of Fv1 produced Fv7, which encodes distinct restriction activity that might provide complementary defenses against retroviruses (Yap et al., 2020). Similarly, in New World monkeys, APOBEC3G genes were repeatedly duplicated through retrotransposition to expand the potential for virus restriction by APOBEC proteins (Yang et al., 2020). In both cases, the major function of the duplicated parental gene is virus inhibition. In contrast, retroCHMP3 proteins originate from genes involved in a core cellular process exploited by viruses to facilitate replication. RetroCHMP3 proteins do not interact directly with the viral pathogen but instead target a cellular pathway. This raises the intruiging possibility that retroCHMP3 may be relatively resistant to rapid counteradaptations by viruses.

Another noteworthy difference between retroCHMP3 proteins and other antiviral proteins is the sporadic distribution of intact genes encoding retroCHMP3, even among closely related species. While there are several other examples of dynamic species-specific expansion and loss of restriction factors (Sawyer et al., 2007; Yang et al., 2020; Young et al., 2018), most antiviral proteins are conserved across mammalian species, or even all the way back to the common ancestor of vertebrates (Bernheim et al., 2021; Blanco-Melo et al., 2016; Conticello et al., 2005; Heusinger et al., 2015; Monit et al., 2019; Sawyer et al., 2007). One notable exception is Fv1, which is derived from an endogenous retrovirus and is found only in *Mus* and a few other rodent species (Benit et al., 1997; Best et al., 1996). In contrast, retroCHMP3 genes are distributed among primates and rodents as full-length, truncated, and pseudogenized copies but are not conserved across mammalian species. The large pool of pseudogenized copies of retroCHMP3 (**Fig. 1B**) is likely to include variants that provided antiviral activity in ancestors of modern species but were subsequently lost, which highlights the cost/benefit tradeoff of altering an essential cellular process like the ESCRT pathway.

The striking diversity of retroCHMP3 variants suggests a flexible mechanism for repeated generation of potent immune defenses. Even from our relatively small sampling of primates and rodents, we discovered a collection of independently truncated retroCHMP3 genes that may provide active antiviral defenses (**Fig. 1**). Several species of New World monkeys encode full-length variants that are one truncating mutation away from providing potent antiviral activity (**Fig. 2, 4**) highlighting a potential reservoir of immune defenses of the future. The recurrent emergence of retroCHMP3 in diverse species raises the possibility that additional retrogenes derived from genes involved in other essential cell functions hijacked by infectious microbes could turn cellular vulnerabilities into opportunities for potent new immune defenses.

## Acknowledgments

We thank Dustin Hancks, Zoë Hilbert, and Keir Balla for insights and helpful discussion, and Janet Iwasa for artwork. Flow cytometry was performed at the University of Utah Flow Cytometry Core. ddPCR was performed at the Molecular Diagnostics Section of the Biorepository and Molecular Pathology Shared Resource at the Huntsman Cancer Institute, University of Utah. This work was supported by NIH grants R37 AI 51174 (to W.I.S), P50 AI150464 and R35 GM134936 (to N.C.E) and a Burroughs Wellcome Fund Investigators in the Pathogenesis of Infectious Disease award (to N.C.E).

## Author contributions

Conceptualization: D.M.D, L.R, K.A.D, W.I.S, N.C.E. Investigation: L.R, K.A.D, D.M.D, A.N.M. Supervision: W.I.S, N.C.E. Writing – Original Draft: L.R, K.A.D, N.C.E. Writing – Review and Editing: All authors.

## Competing interests

The authors declare no competing interests.

## Supplemental Information

**Fig. S1:**
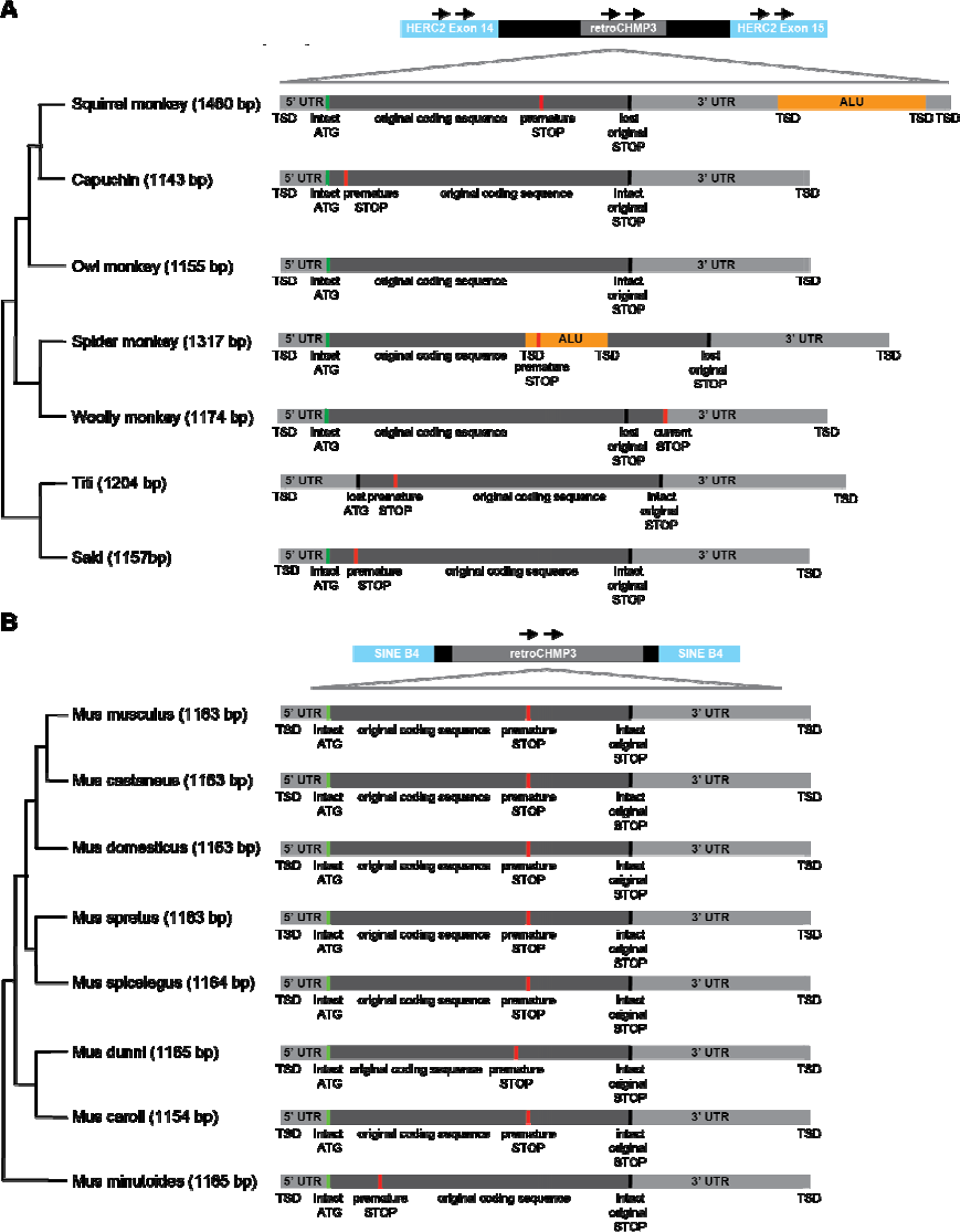
Schematic representation of the syntenic retroCHMP3 genes in New World monkeys (A) and mice (B). Number after species name indicates length of retrotransposed region. TSD – target site duplication. UTR – untranslated region

**Fig. S2:**
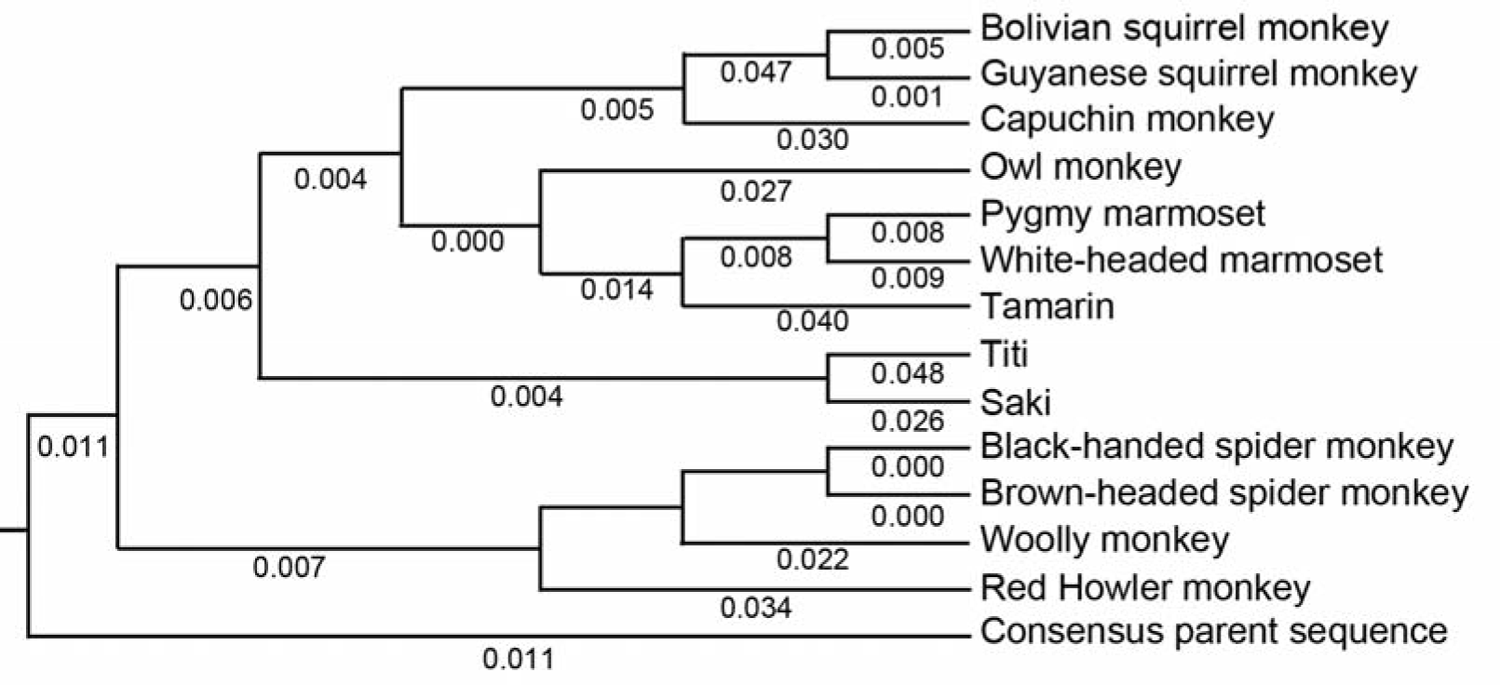
Ancestral reconstruction of New World monkey ancestral retroCHMP3. Parental CHMP3 sequences were aligned, and the highest quality consensus sequence resulting from the alignment was used as an approximation of the ancestral CHMP3 retrocopy (consensus parent). The consensus parent and retroCHMP3 sequences were aligned and used to infer the maximum likelihood phylogeny using the GTR+GAMMA model of rate heterogeneity. The consensus parent sequence was specified as the outgroup. Black numbers indicate substitutions per site.

**Fig. S3:**
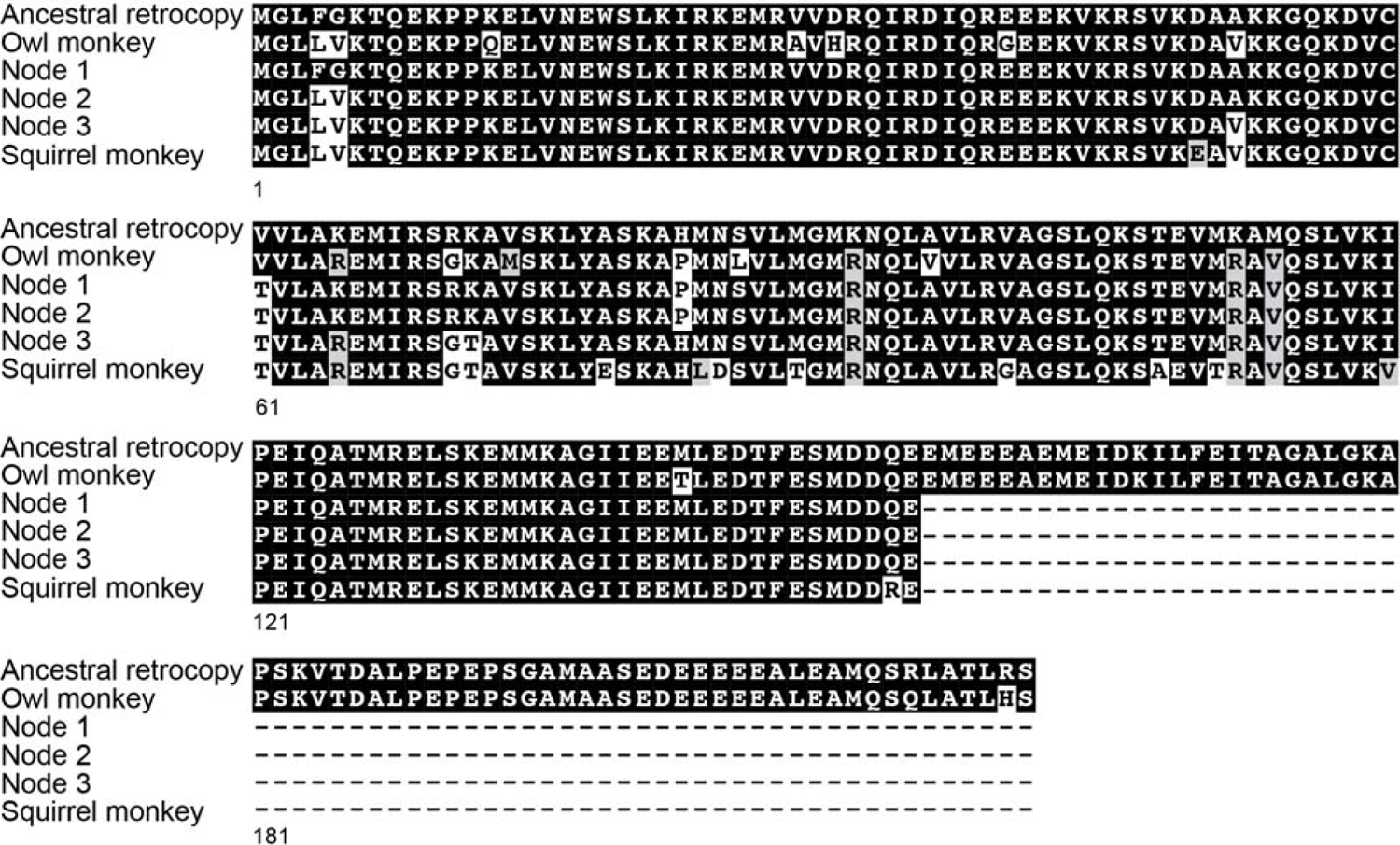
Alignment of reconstructed ancestral retroCHMP3, modern owl monkey retroCHMP3, nodes and modern squirrel monkey retroCHMP3 protein sequences.

**Figure S4:**
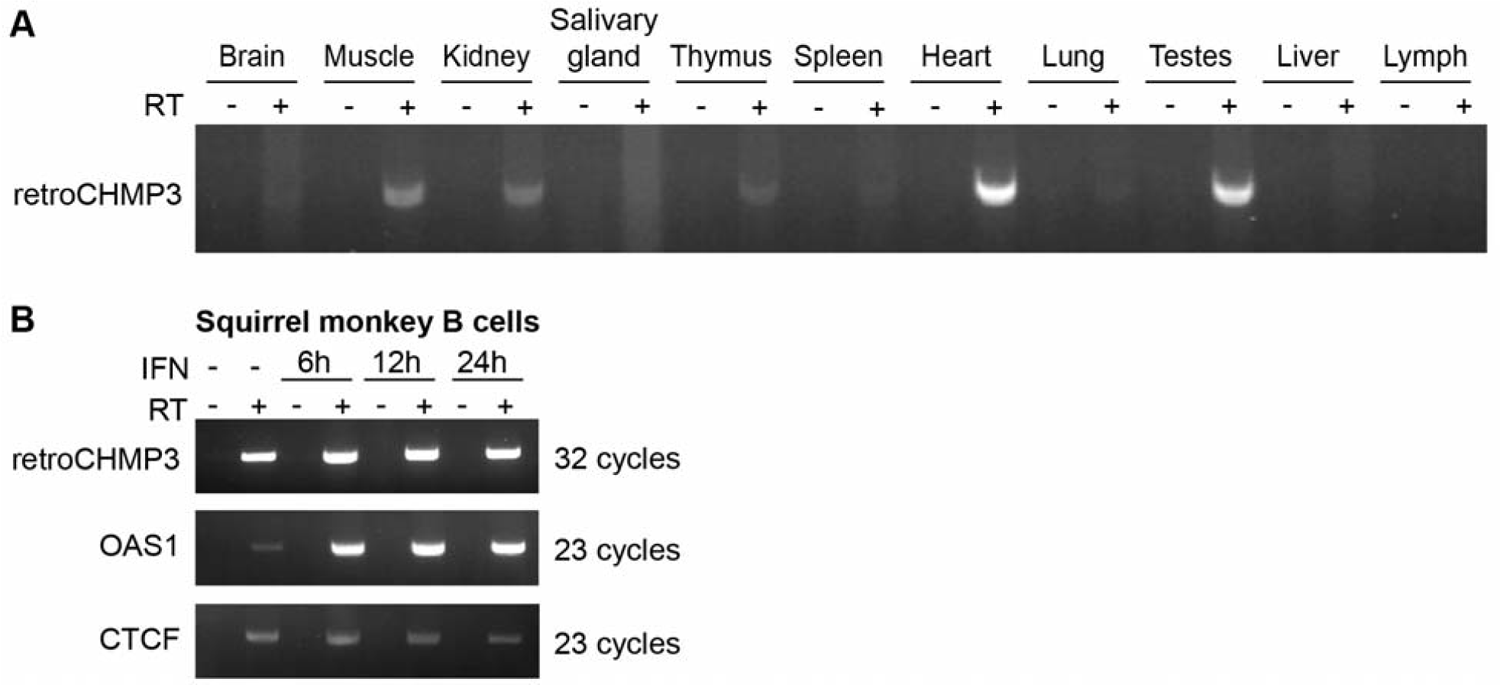
(A) Detection of retroCHMP3 RNA in mouse tissues. **(B)** Detection of retroCHMP3 RNA in squirrel monkey B cells with and without interferon (IFN) treatment. RT – reverse transcriptase. RetroCHMP3 bands were excised, cloned and sequenced to verify a complete match with the retroCHMP3 sequence.

**Table S1:** RetroCHMP3 sequences identified primates Tables show full-length and truncated retrocopies identified in this study. Highly degenerate retrocopies are not included.

**Table S2:** RetroCHMP3 sequences identified in rodents Tables show full-length and truncated retrocopies identified in this study. Highly degenerate retrocopies are not included.

**Table S3:** **Retrocopies of CHMP3, CHMP2A and CHMP4B in primate and rodent genomes.** Species were analyzed for the presence of at least one retrocopy of CHMP3, CHMP2A or CHMP4B by NCBI BLAST of whole-genome shotgun contigs and data were collapsed into families. Numbers indicated number of retrocopies per family. Degenerate retrocopies are not included. *multiple copies within species, **includes woolly monkey and pygmy marmoset retroCHMP3, which were identified by sequencing of genomic DNA.

| Primates (# species investigated) | CHMP3 |  | CHMP2A |  | CHMP4B |  |
| --- | --- | --- | --- | --- | --- | --- |
|  | 140 – 190 aa | Full-length | 140 – 190 aa | Full-length | 140 – 190 aa | Full-length |
| **Hominoidea (9)** | 0 | 0 | 0 | 1 | 0 | 0 |
| **Cercopithecidae (21)** | 2 | 1 | 0 | 3 | 0 | 0 |
| **Platyrrhini (11)** | 2 | 3** | 0 | 1 | 0 | 0 |
| **Tarsiiformes (1)** | 0 | 0 | 0 | 0 | 0 | 0 |
| **Lorisiformes (2)** | 1 | 2 | 0 | 0 | 0 | 0 |
| **Lemuridae (5)** | 0 | 1 | 0 | 0 | 0 | 0 |
| **Cheirogaleus (1)** | 2* | 0 | 0 | 0 | 0 | 0 |
| **Mirza (2)** | 0 | 5* | 0 | 0 | 0 | 0 |
| **Microcebus (6)** | 1 | 5 | 0 | 0 | 0 | 0 |
| Rodents (# species investigated) | CHMP3 |  | CHMP2A |  | CHMP4B |  |
|  | 140 – 190 aa | Full-length | 140 – 190 aa | Full-length | 140 – 190 aa | Full-length |
| **Mus (7)** | 6 | 0 | 0 | 0 | 0 | 0 |
| **Grammomys (1)** | 0 | 1 | 0 | 0 | 0 | 0 |
| **Rattus (2)** | 0 | 0 | 0 | 0 | 0 | 1 |
| **Peromyscus (9)** | 5 | 1 | 0 | 0 | 0 | 0 |

**Table S4:**
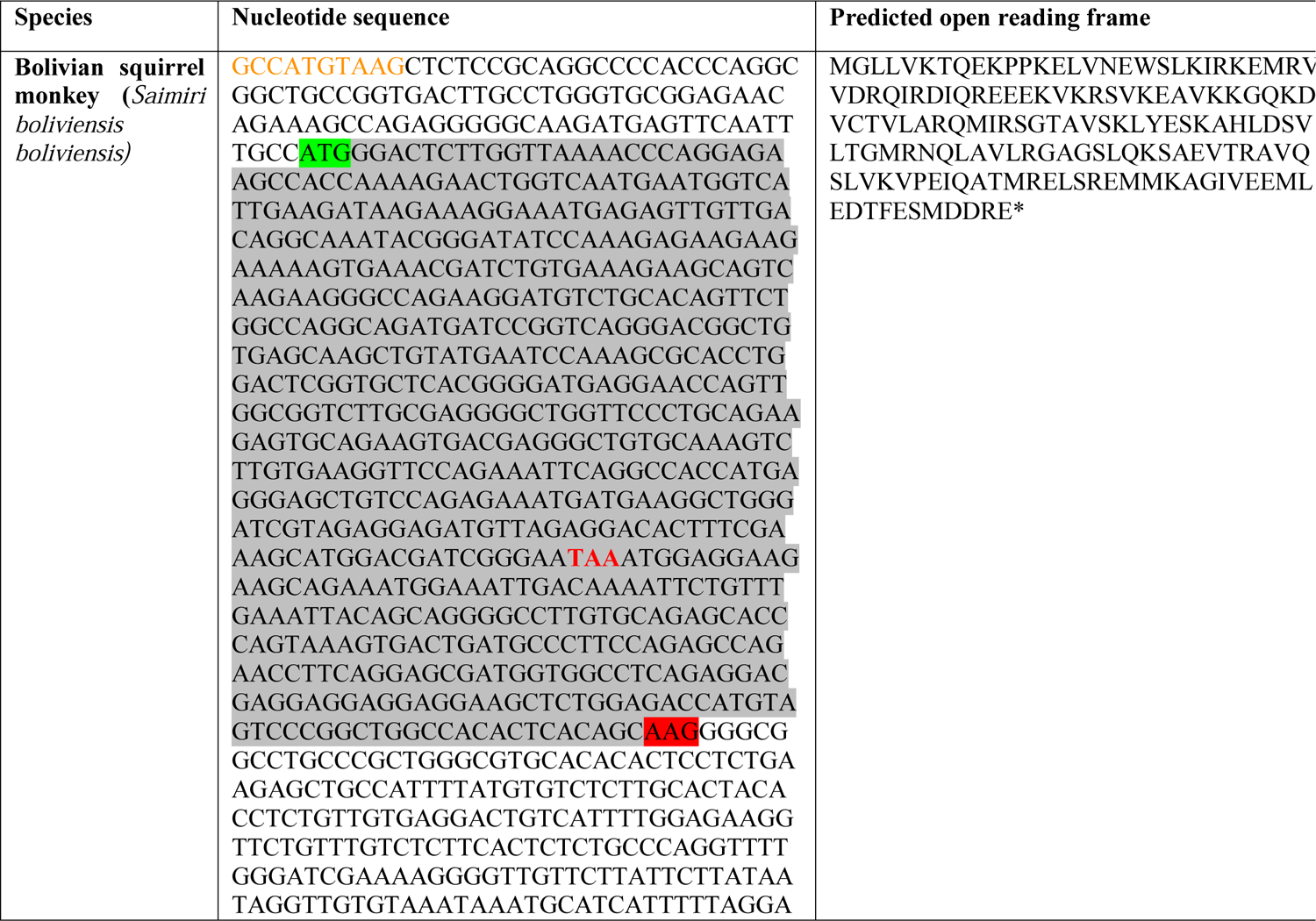
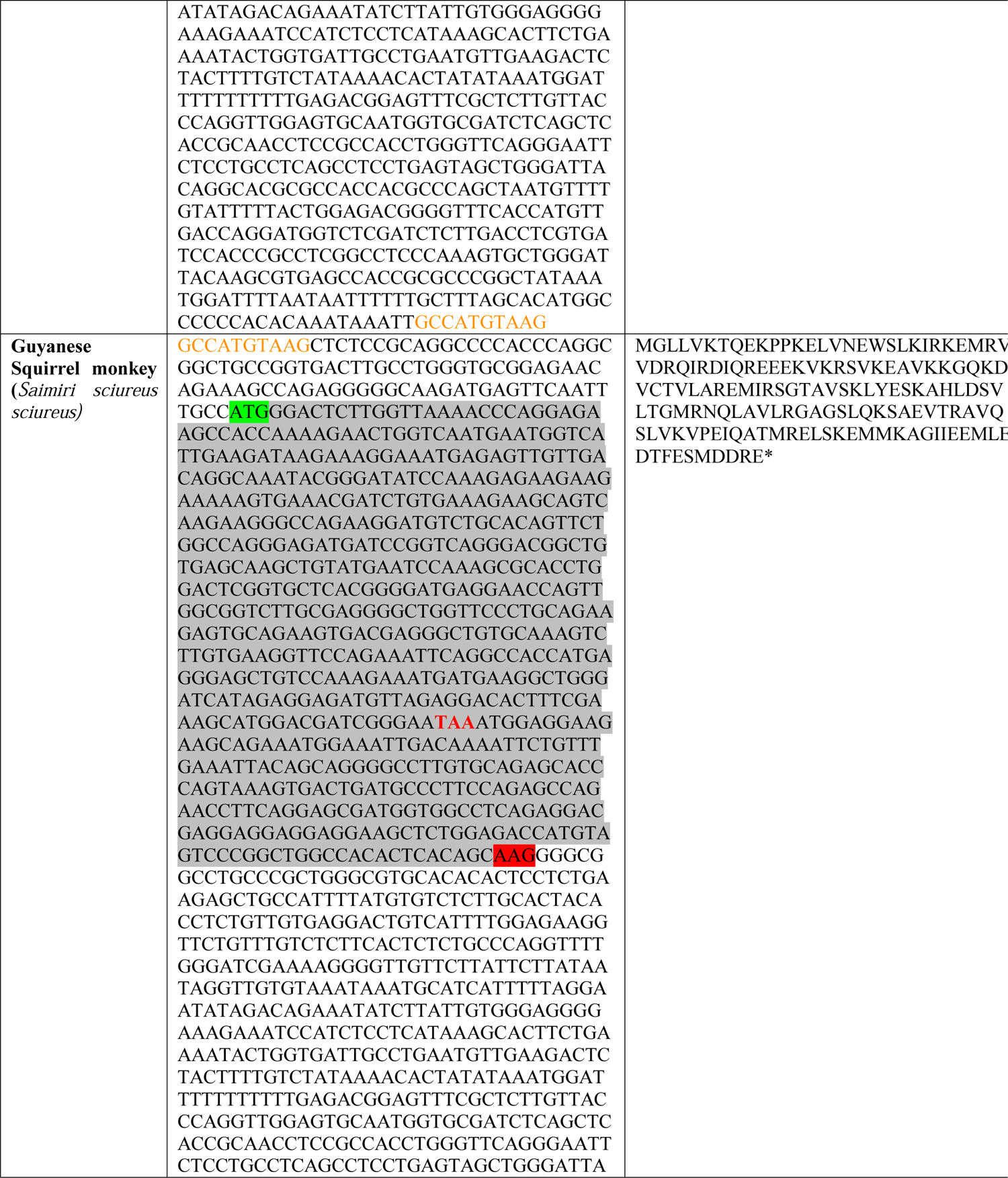
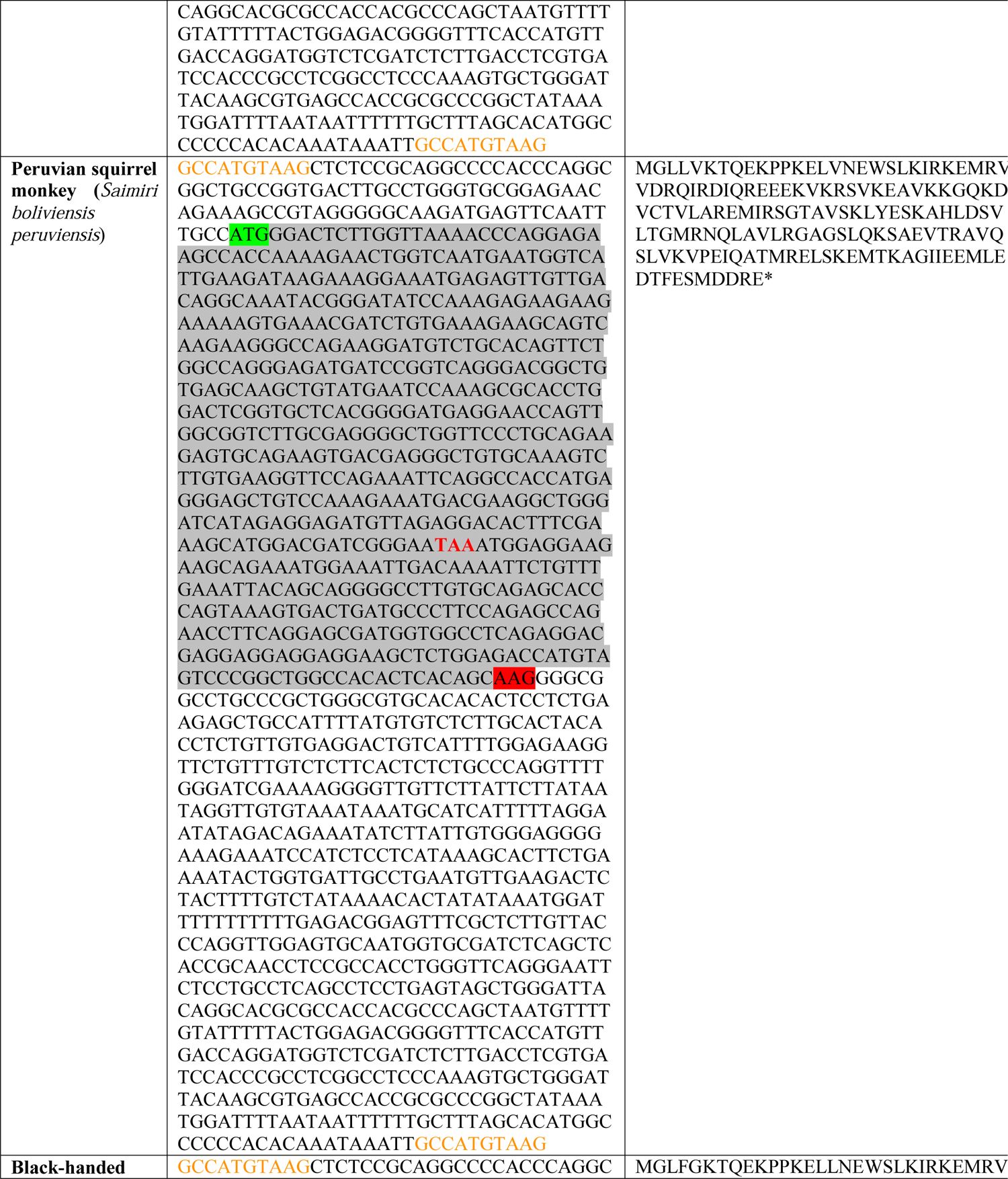
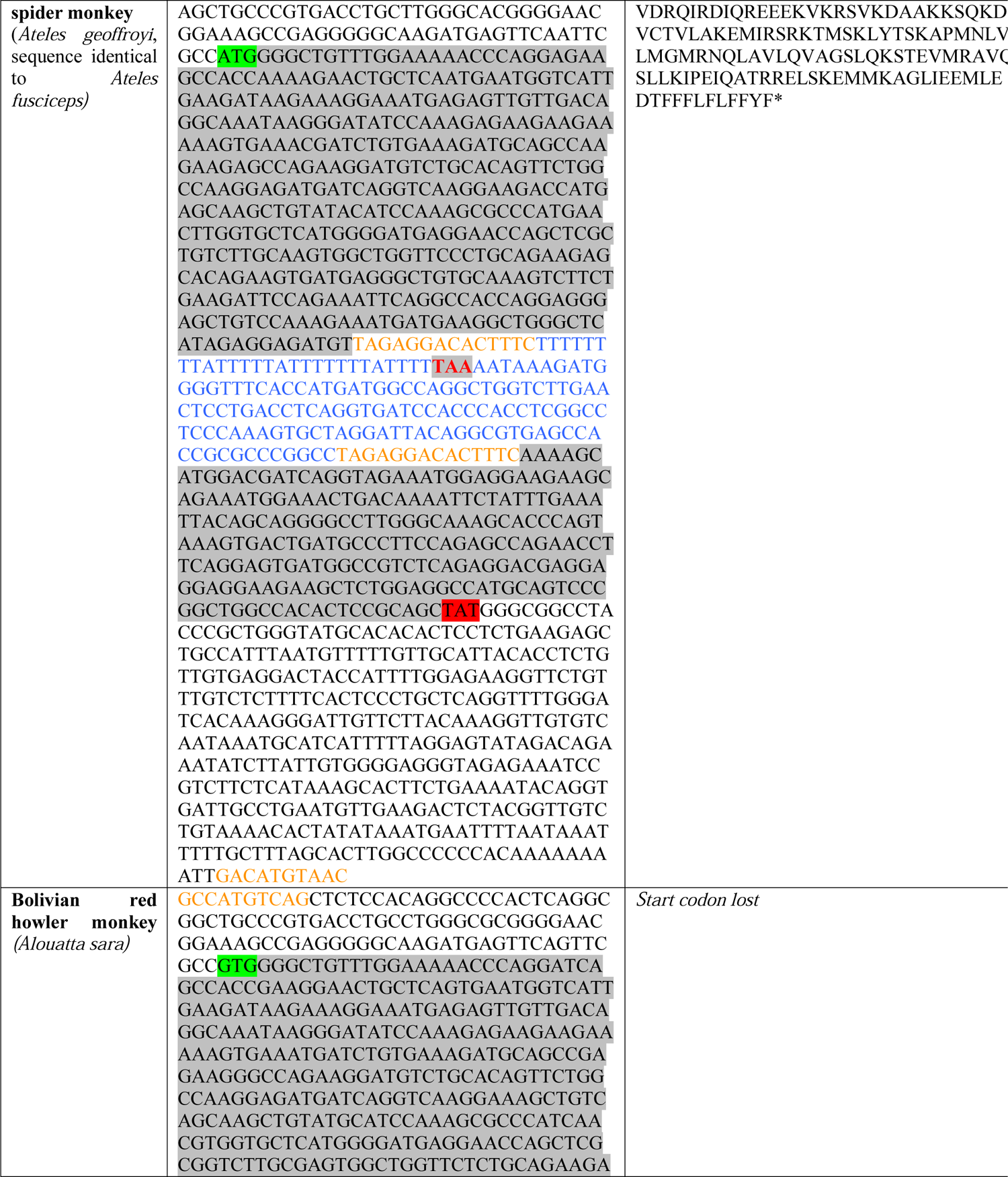
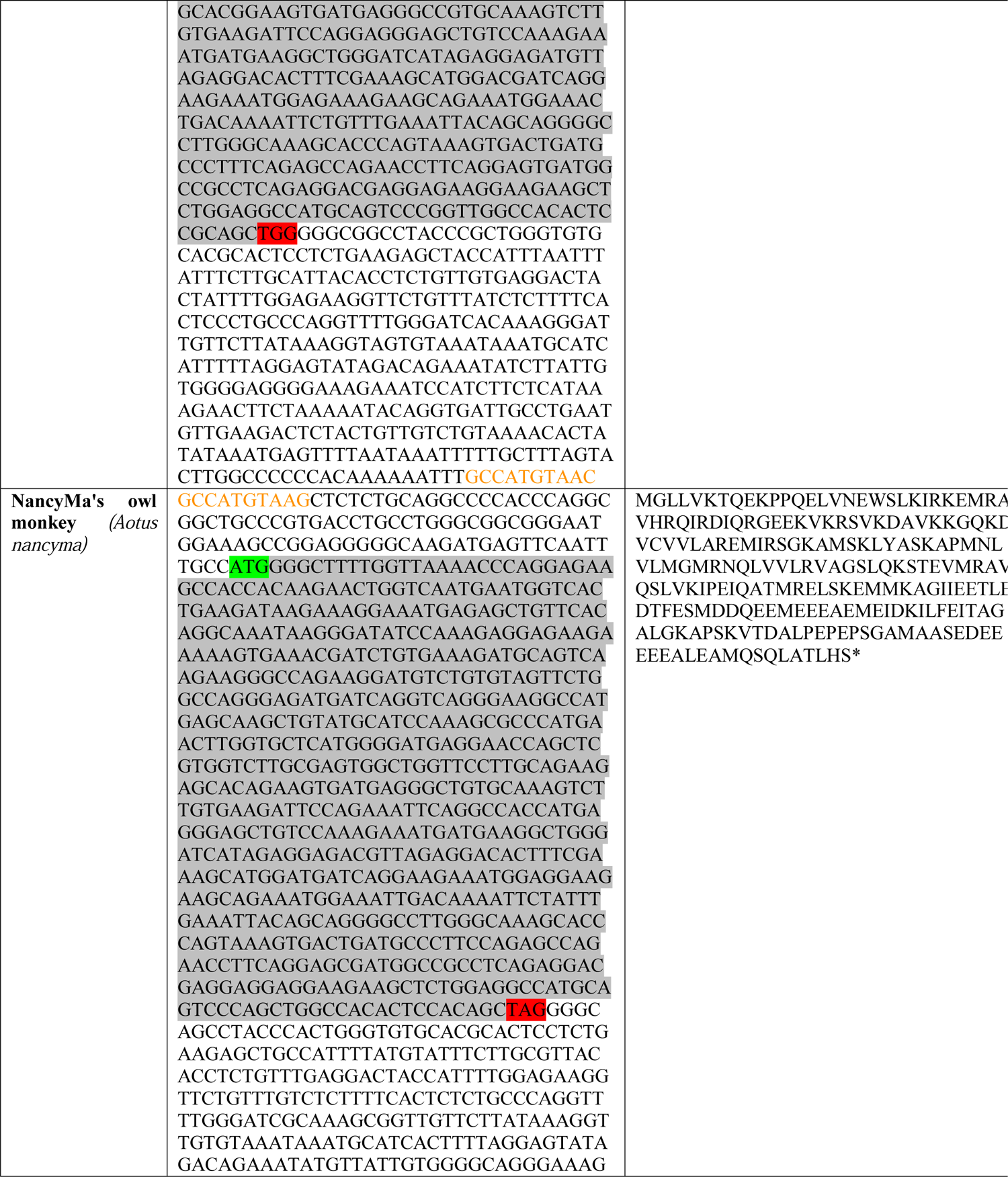
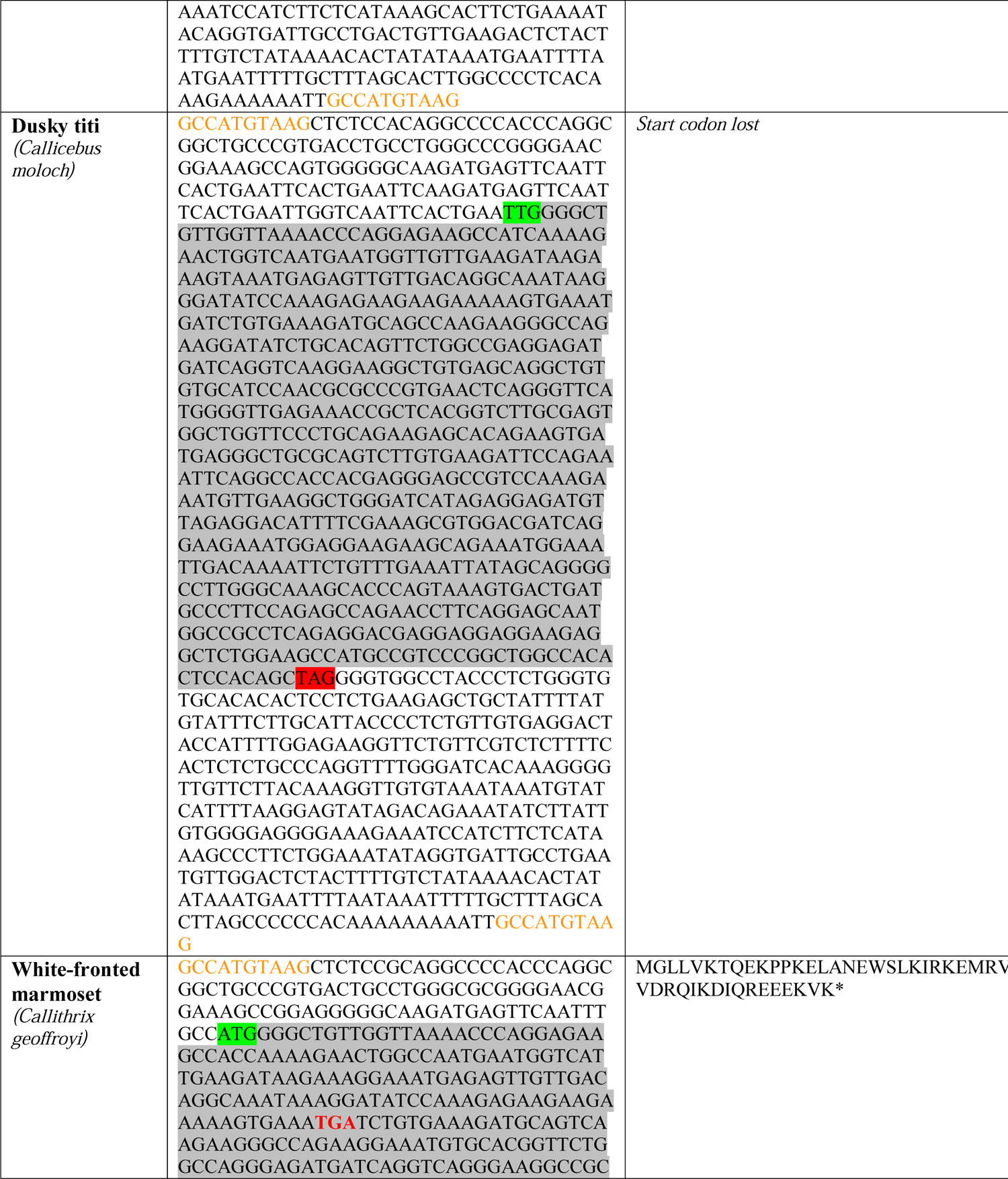
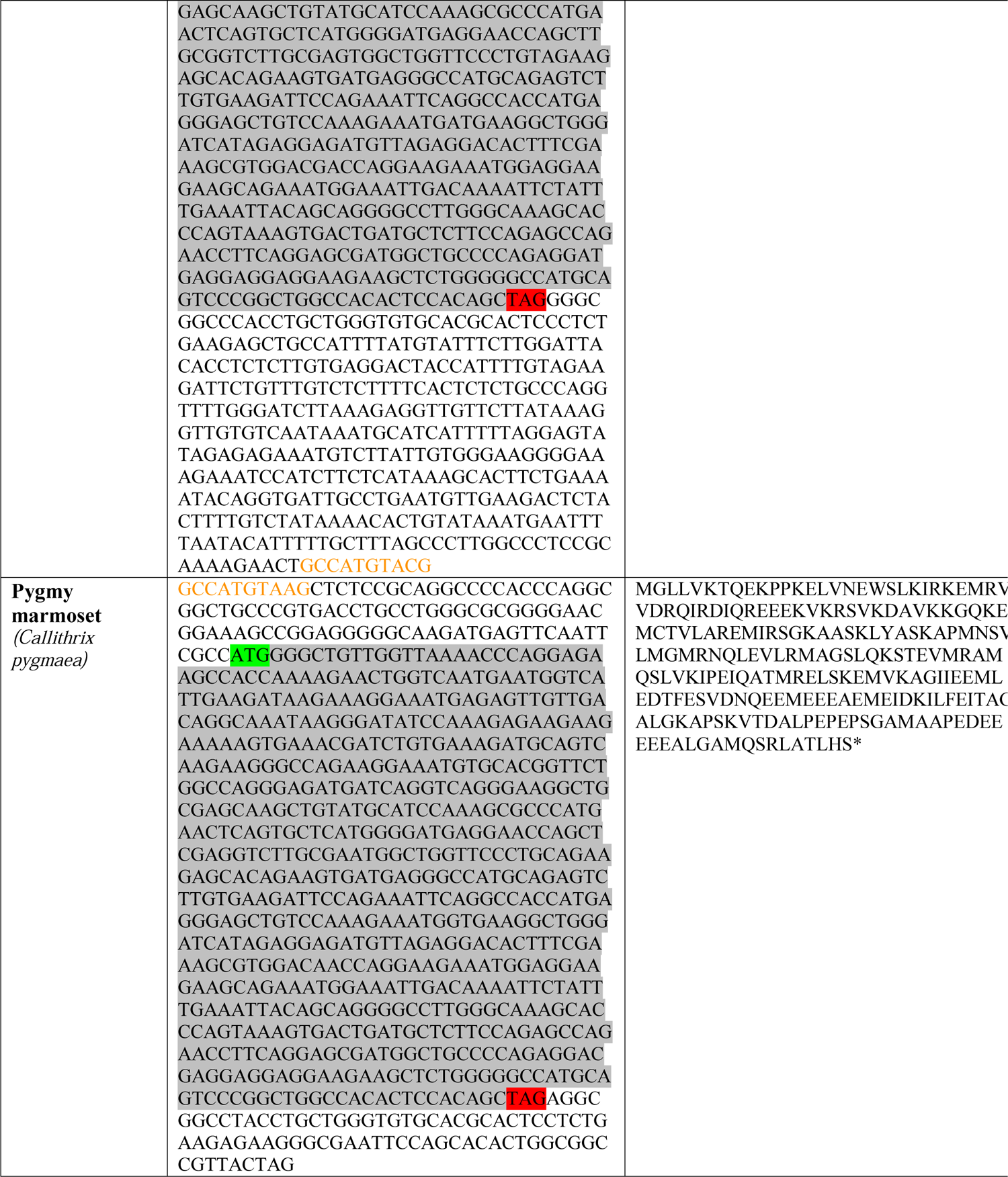
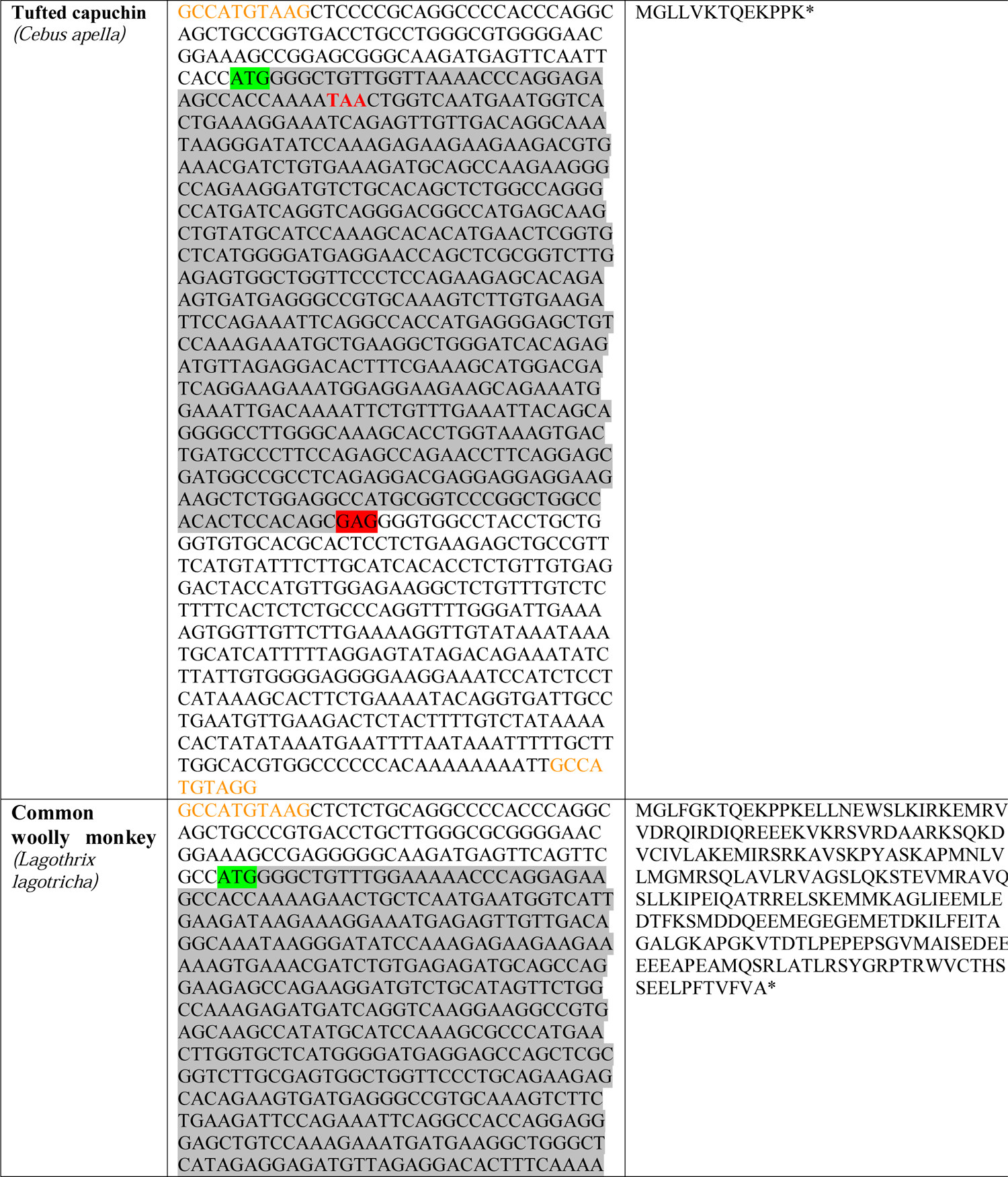
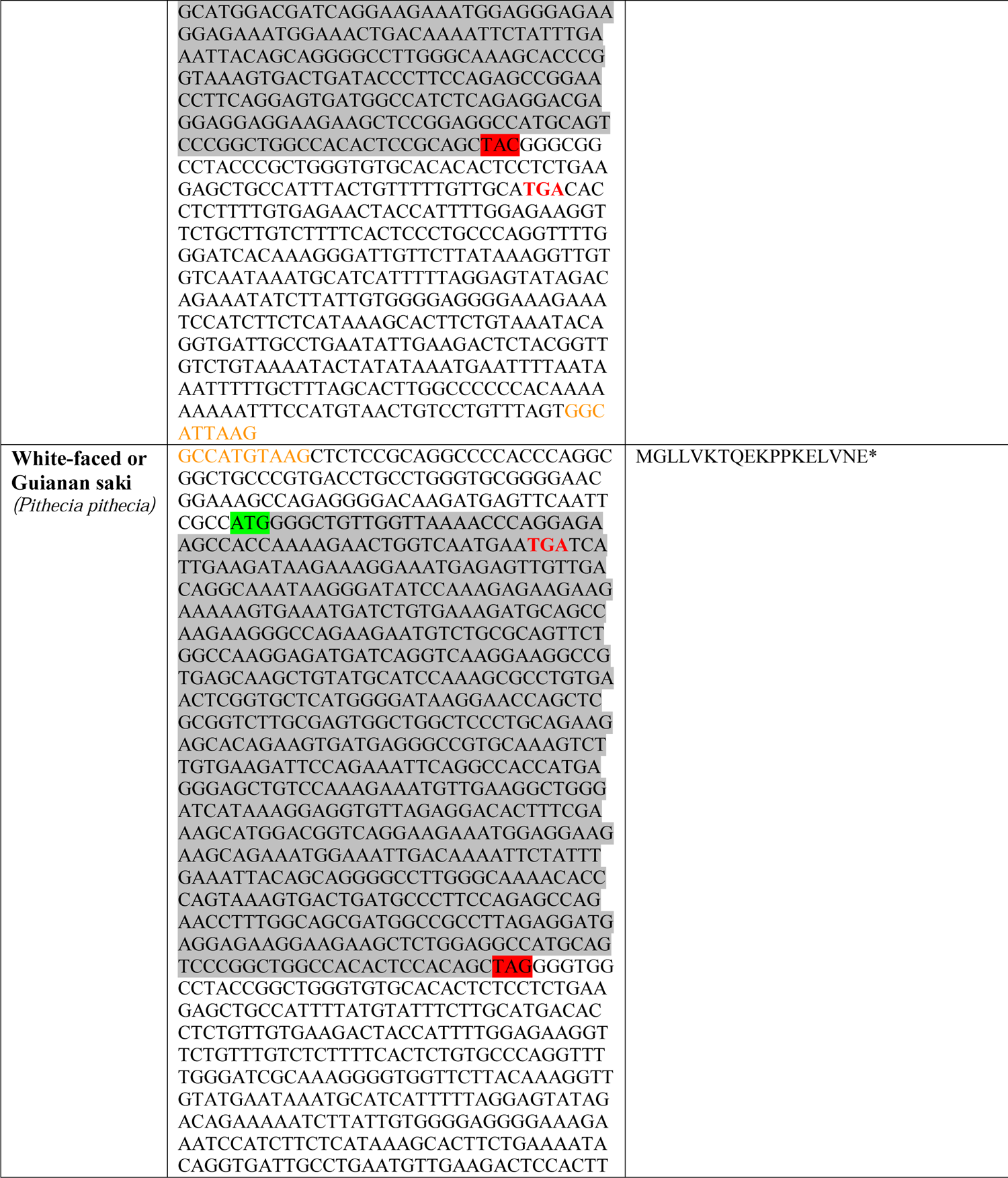
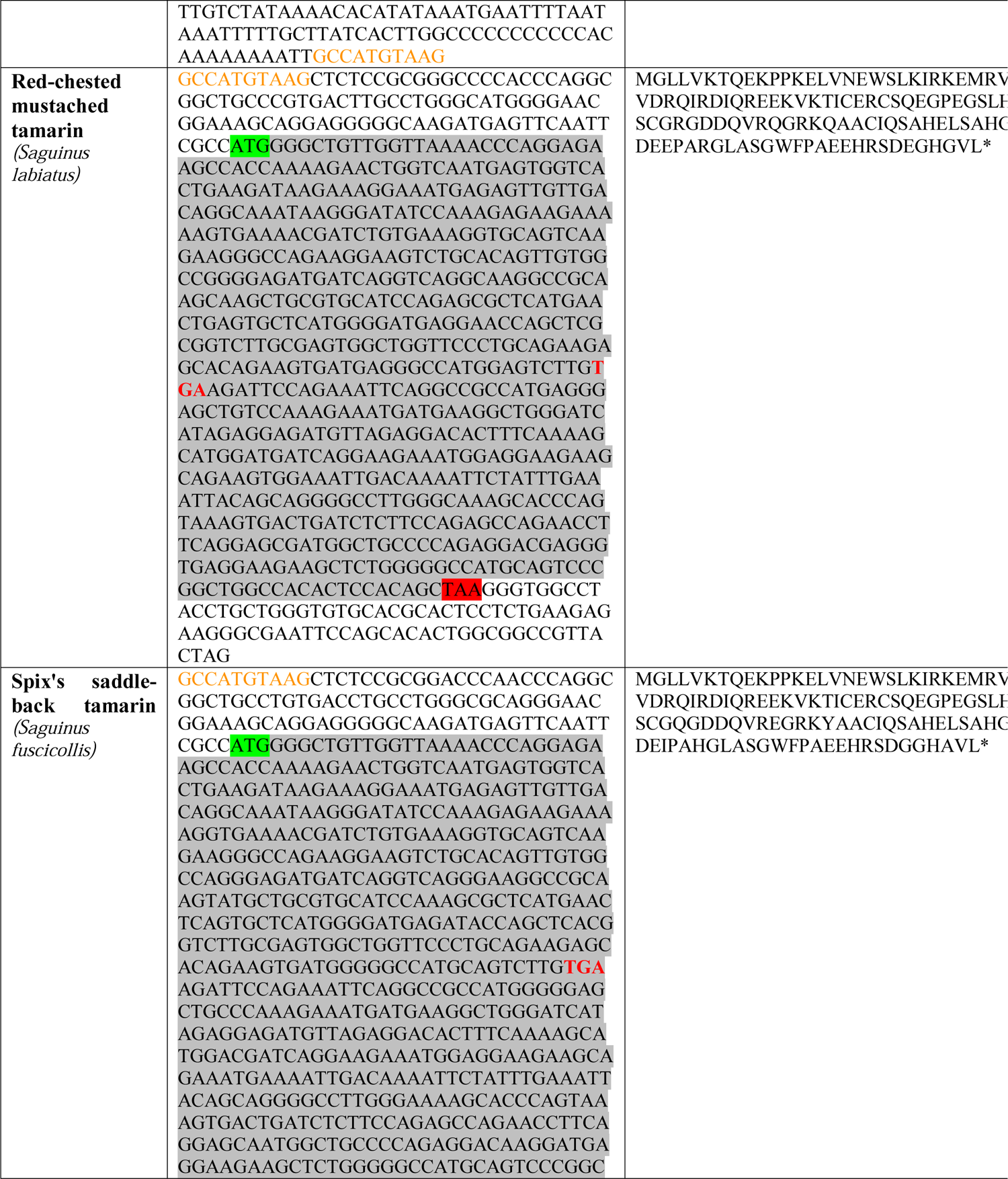
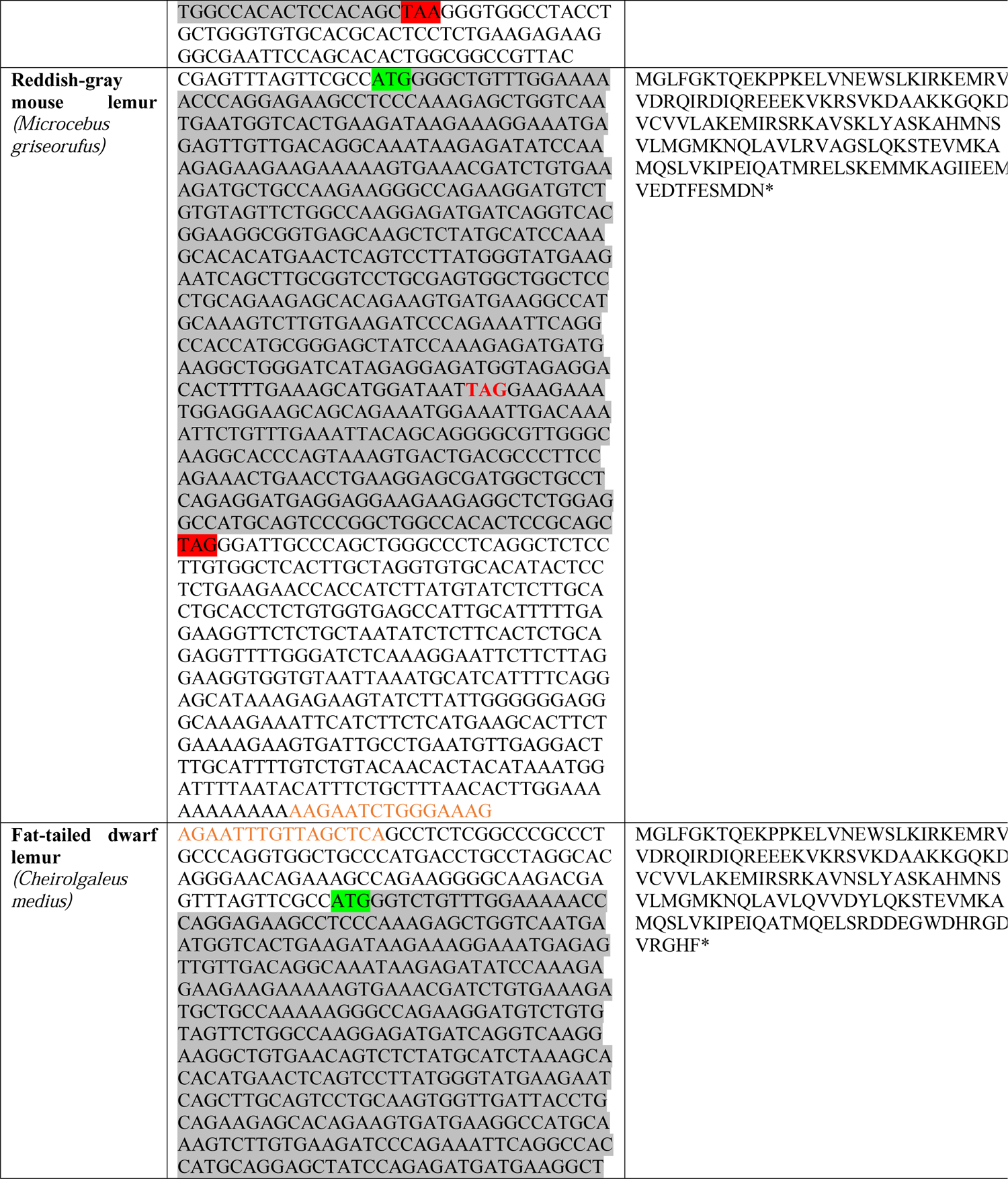
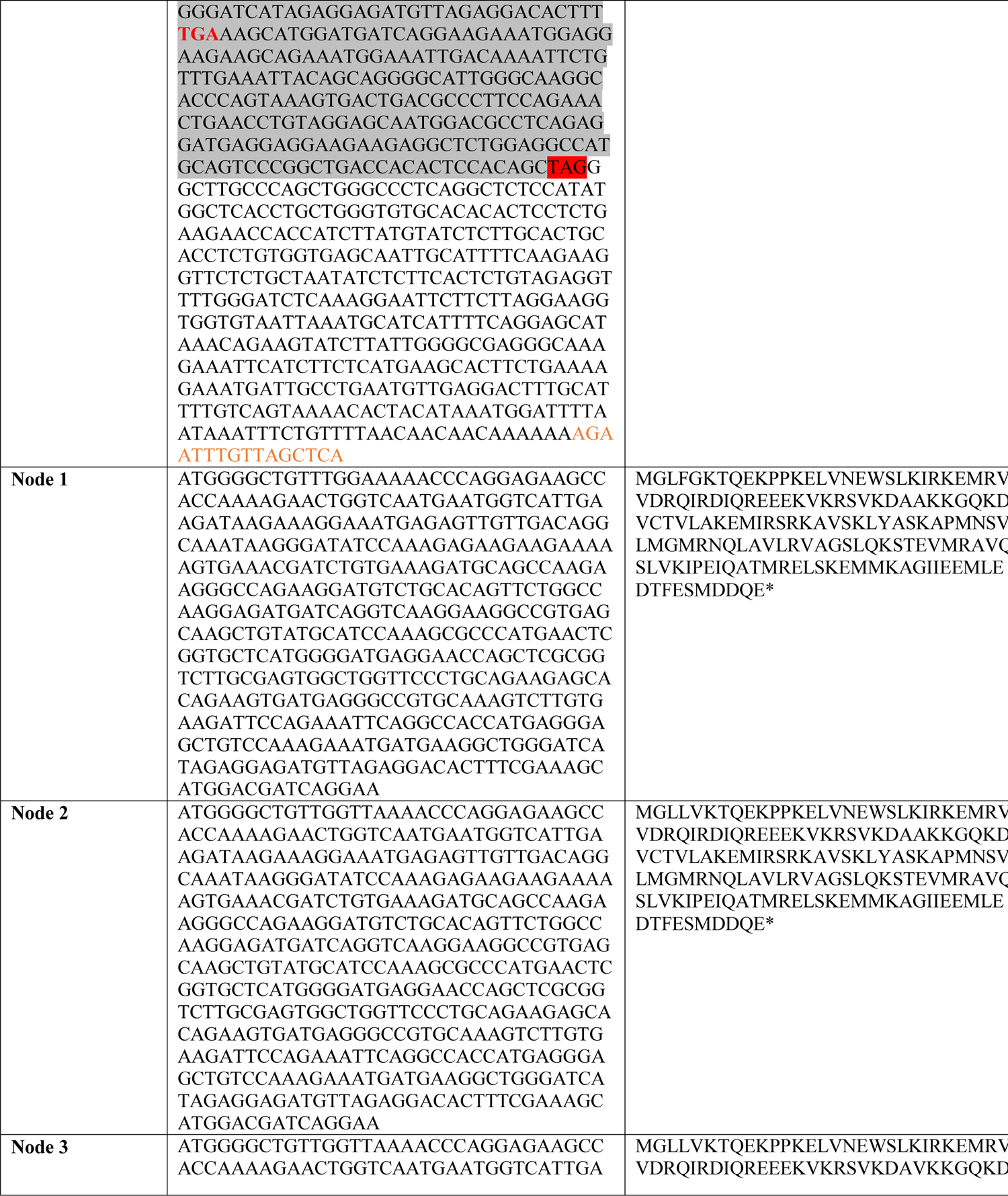
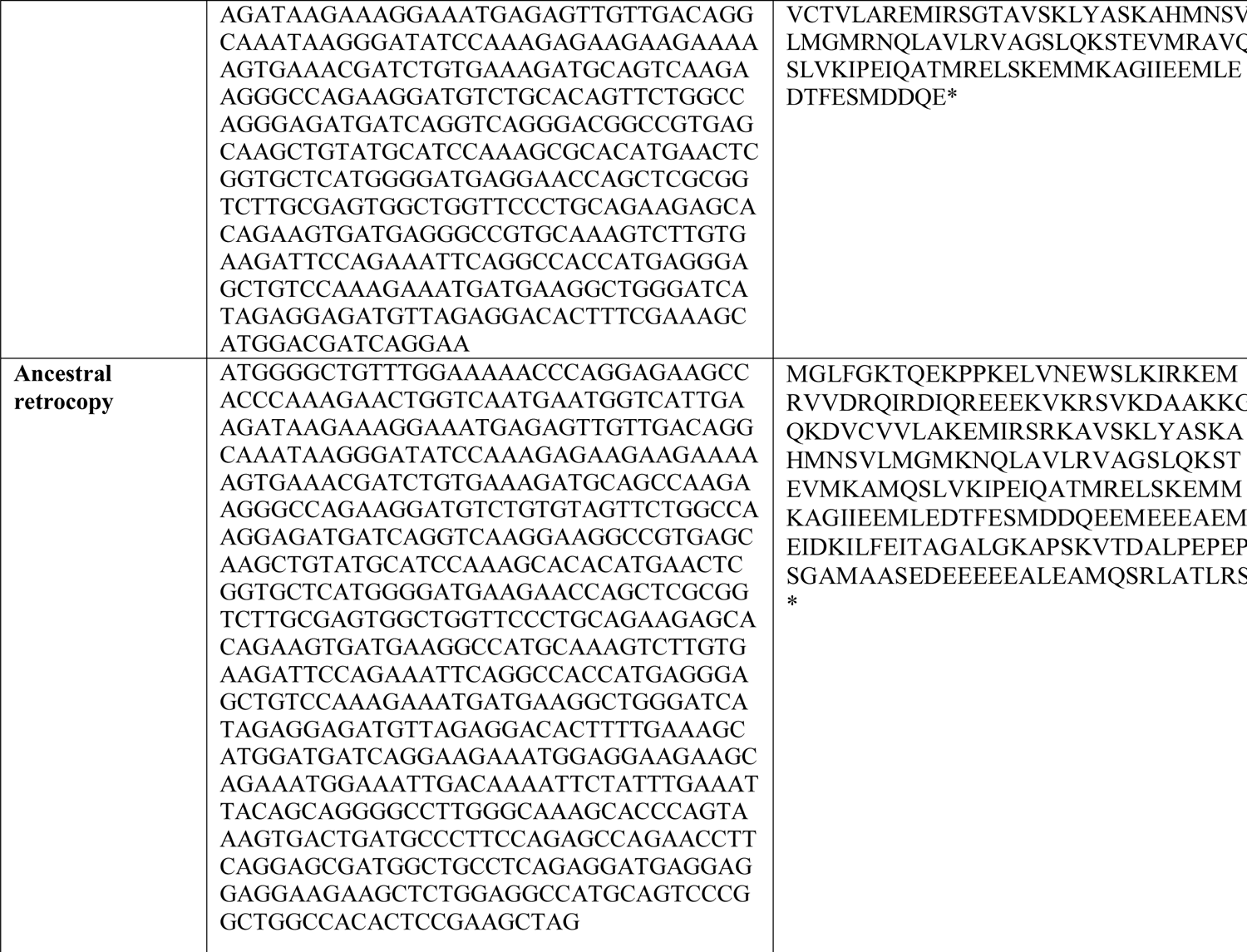
RetroCHMP3 sequences in primates and evolutionary intermediates The color coding and shading highlights key features of each retrocopy. Orange letters indicate TSD sequences flanking the retrocopy. In some cases, there is a single nucleotide difference in the 5’ and 3’ flanking TSD sequences. Gray shading indicates the original CHMP3 coding sequence. Green shading indicates the original start codon, which in some species has been modified and is no longer functional. Red shading indicates the location of the original stop codon. In some cases, this sequence has been modified and is therefore no longer a functional stop. Red TAG or TAA indicates location of premature truncating stop codon, or in Woolly monkey the current predicted stop codon. Blue letters in spider monkey sequence indicate an Alu sequence inserted within the retrocopy coding region, with orange letters indicating the TSD flanking this insertion.

**Table S5:**
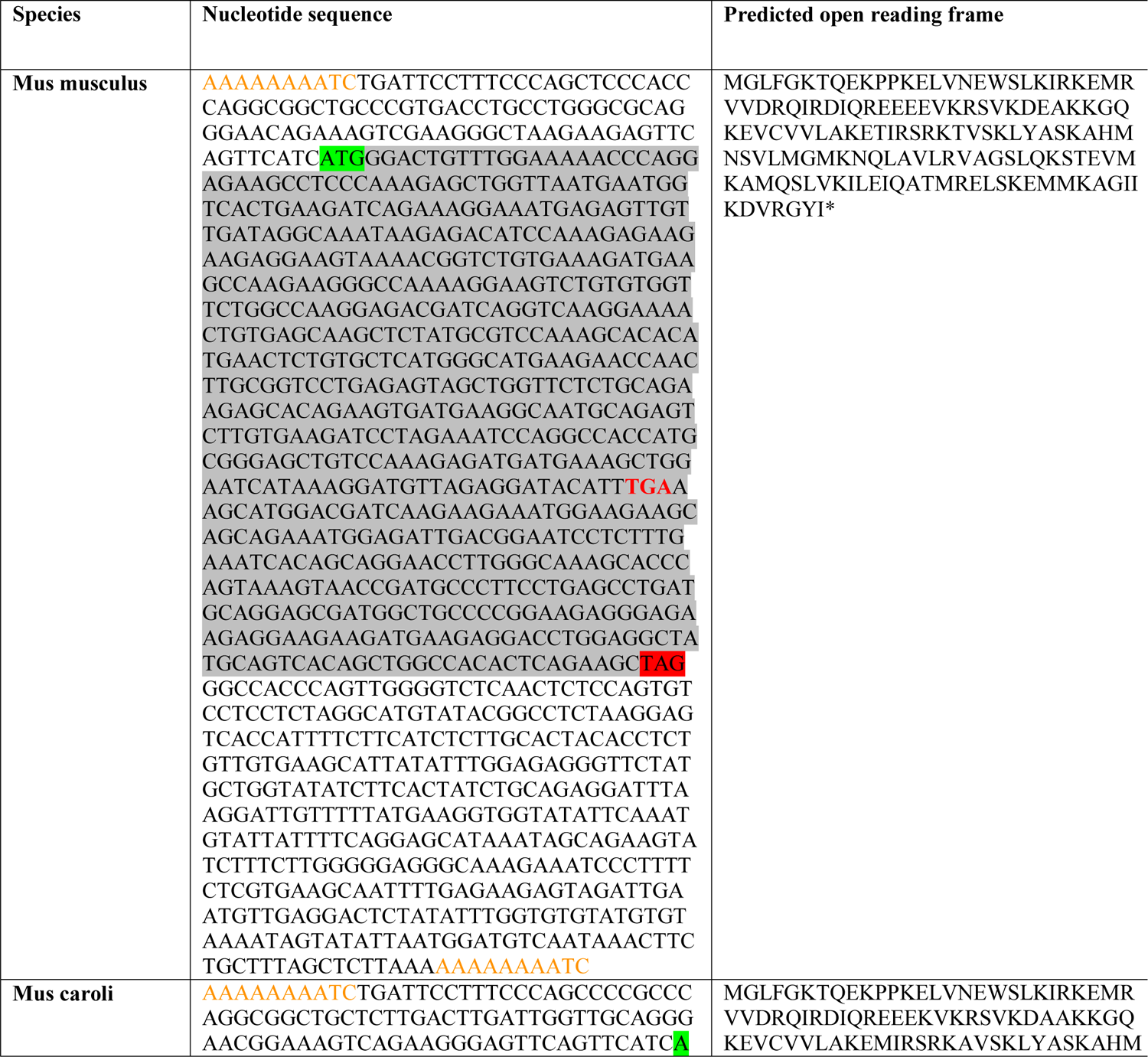
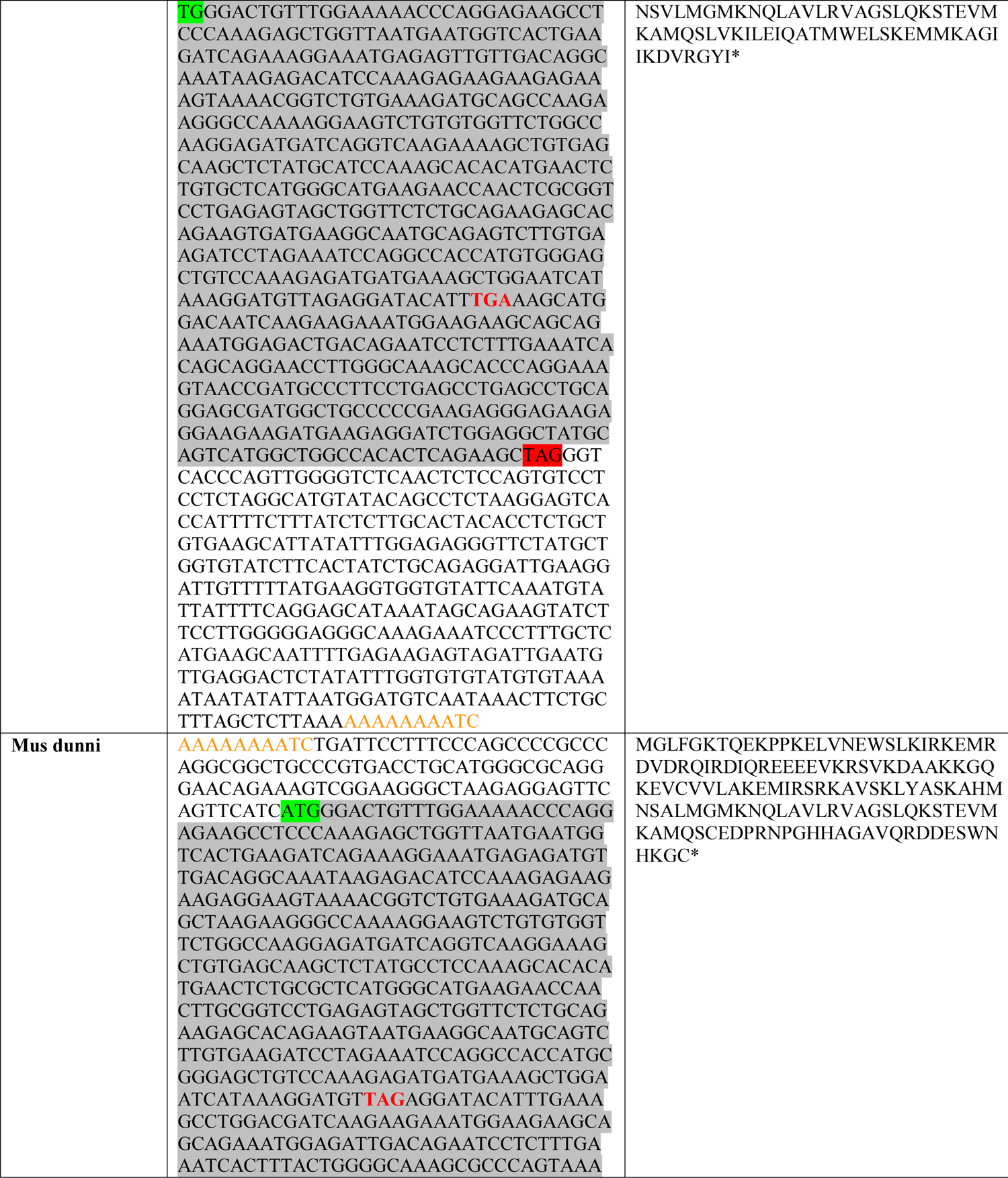
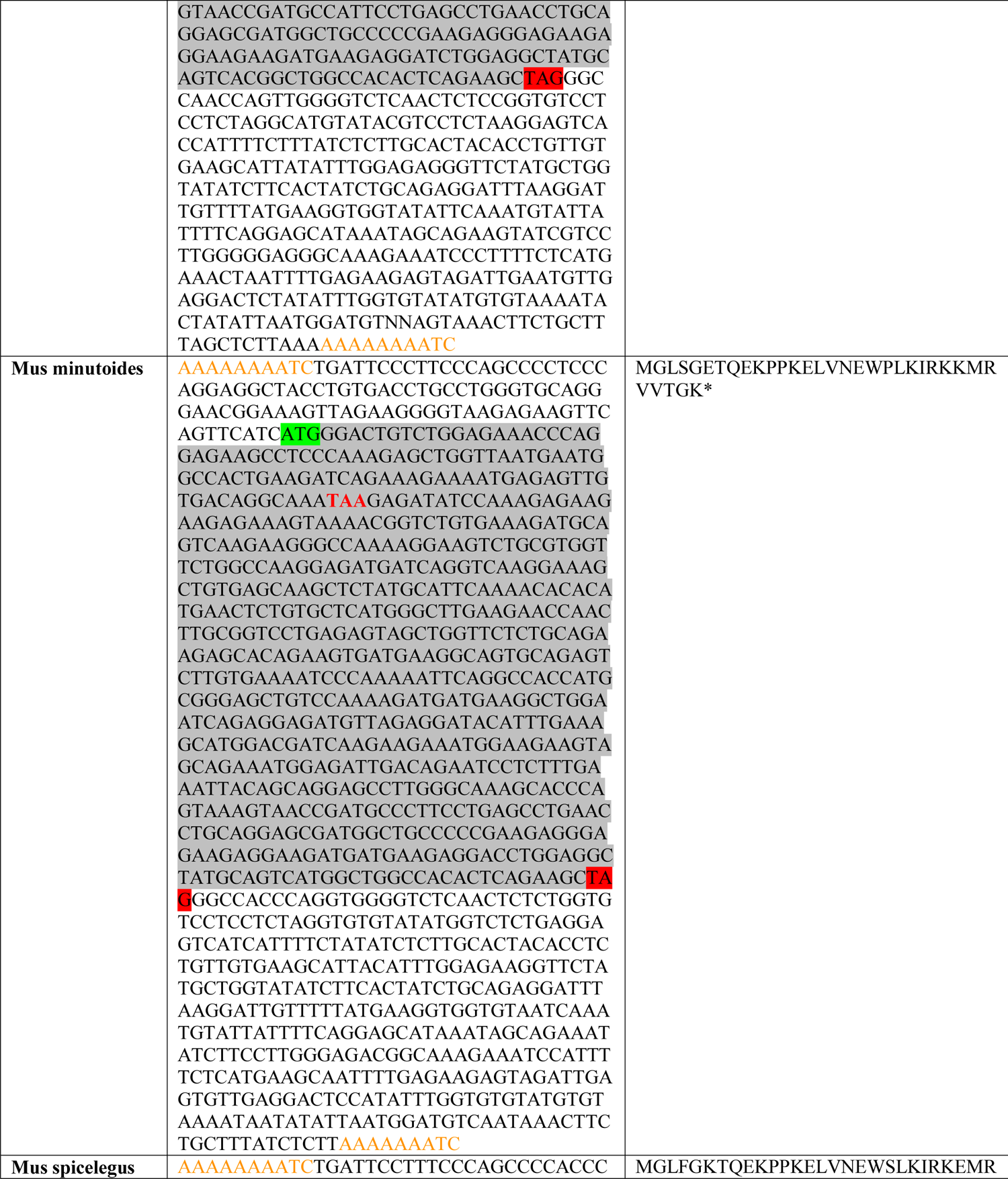
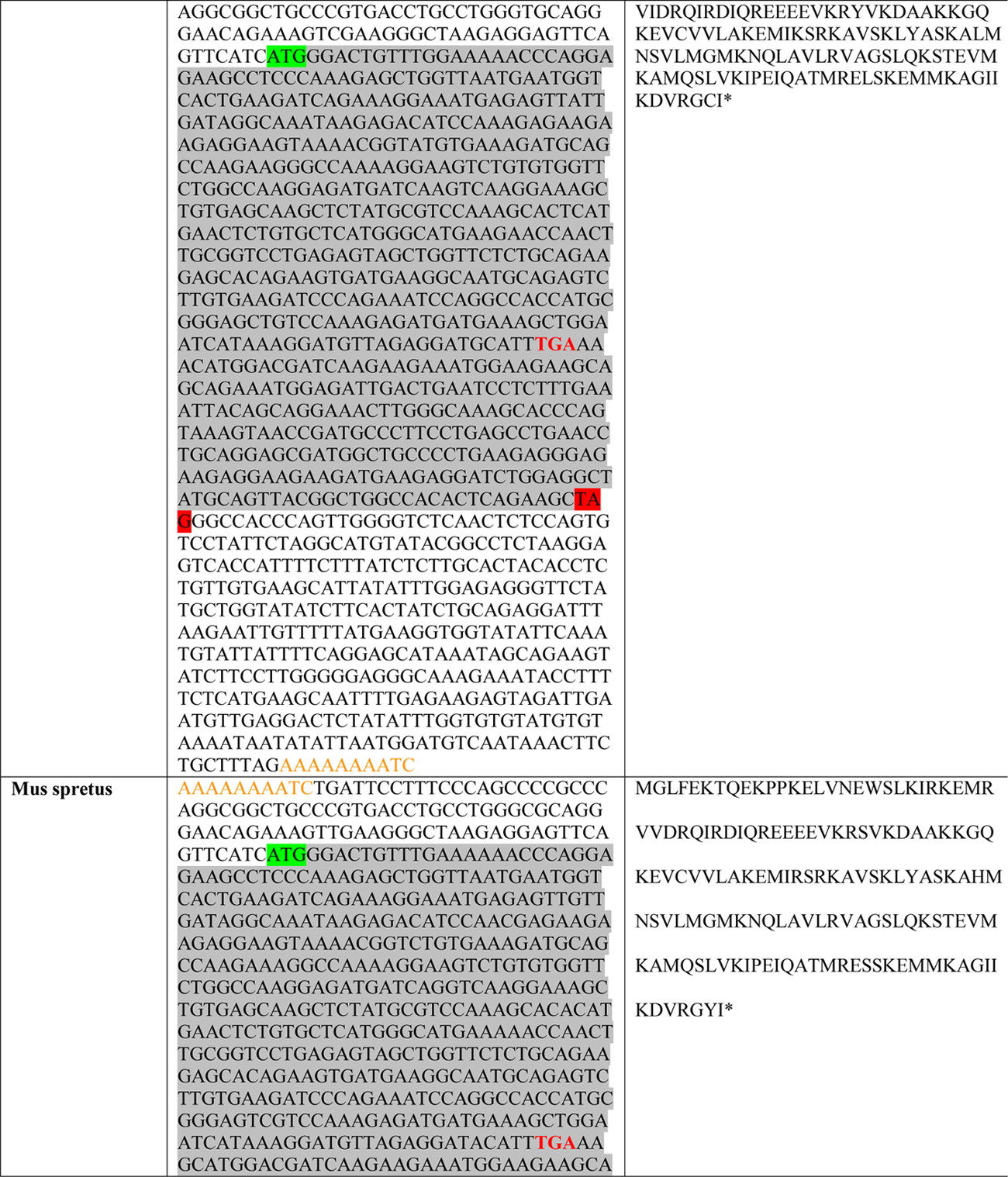
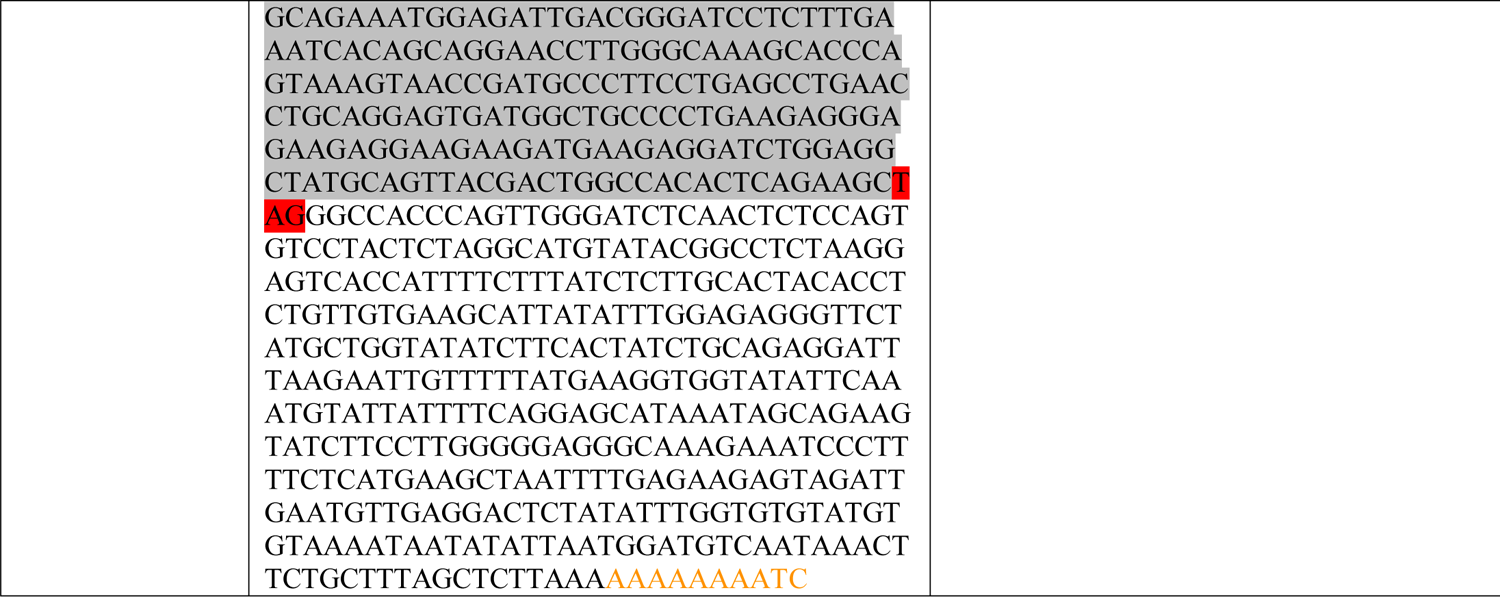
RetroCHMP3 sequences in *Mus* species The color coding and shading highlights key features of each retrocopy. Orange letters indicate TSD sequences flanking the retrocopy. In some cases, there is a single nucleotide difference in the 5’ and 3’ flanking TSD sequences. Gray shading indicates the original CHMP3 coding sequence. Green shading indicates the original start codon. Red shading indicates the location of the original stop codon. Red TAG indicates location of premature, truncating stop codon.

**Table S6:**
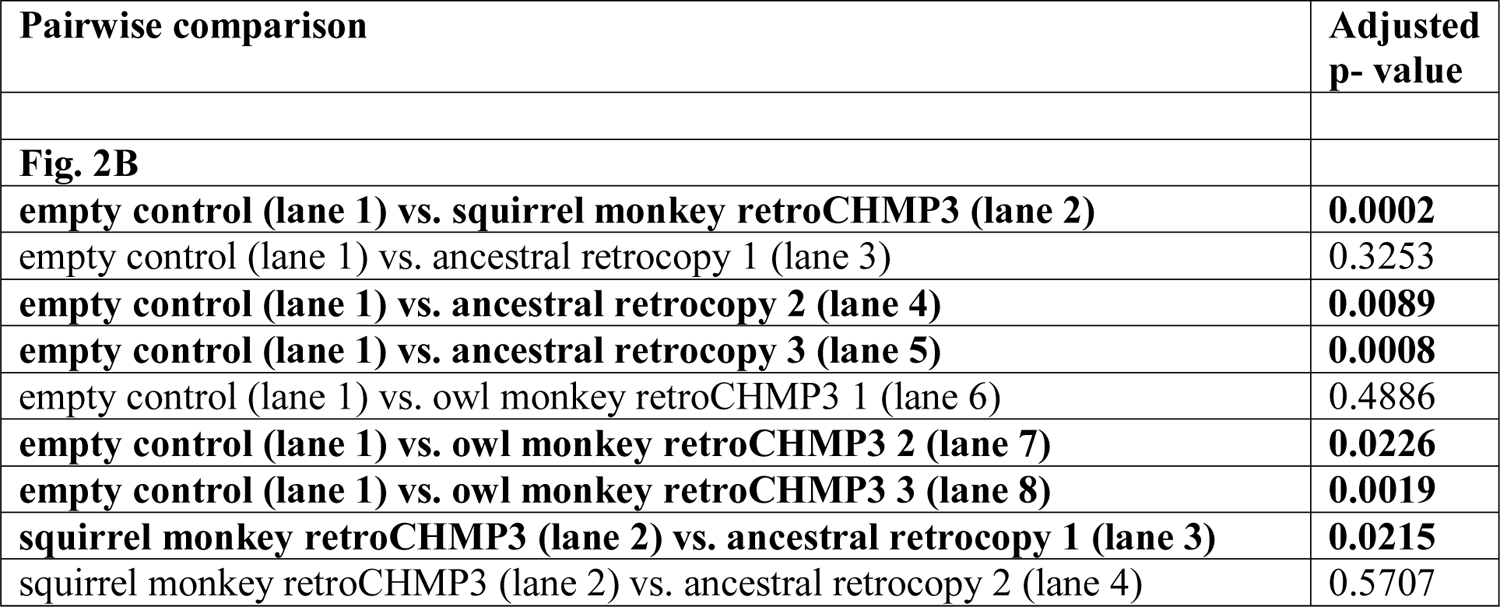
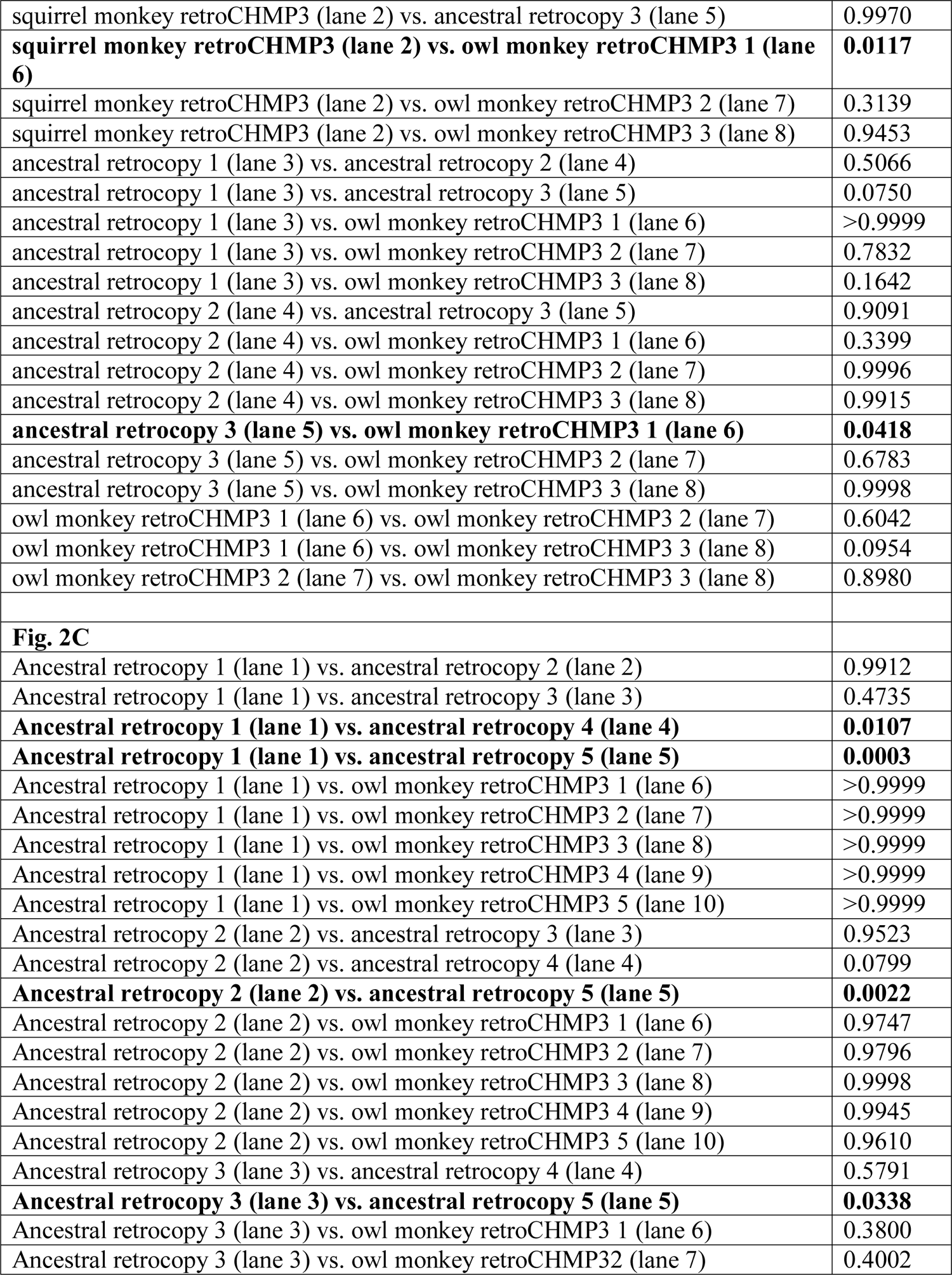
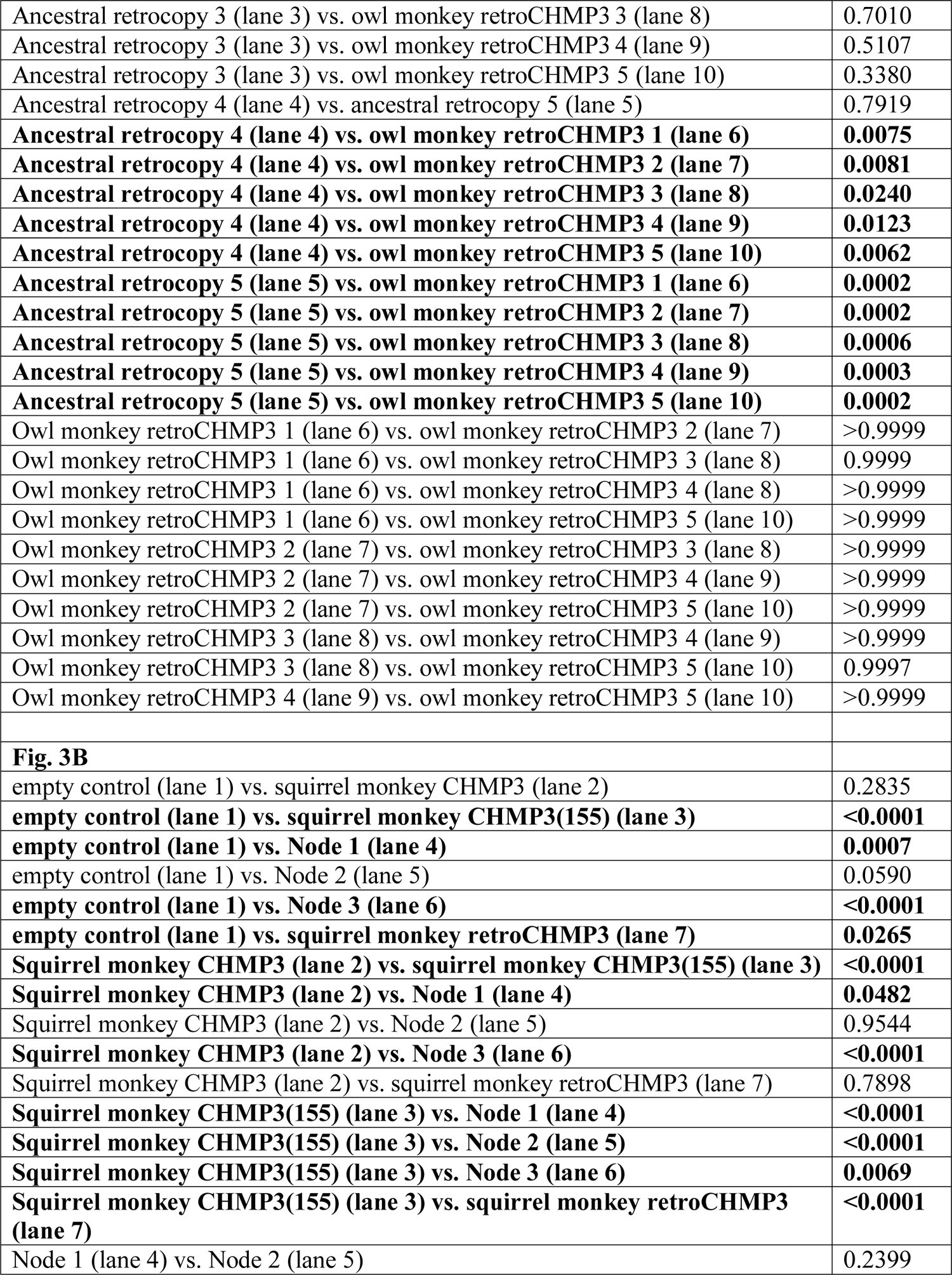
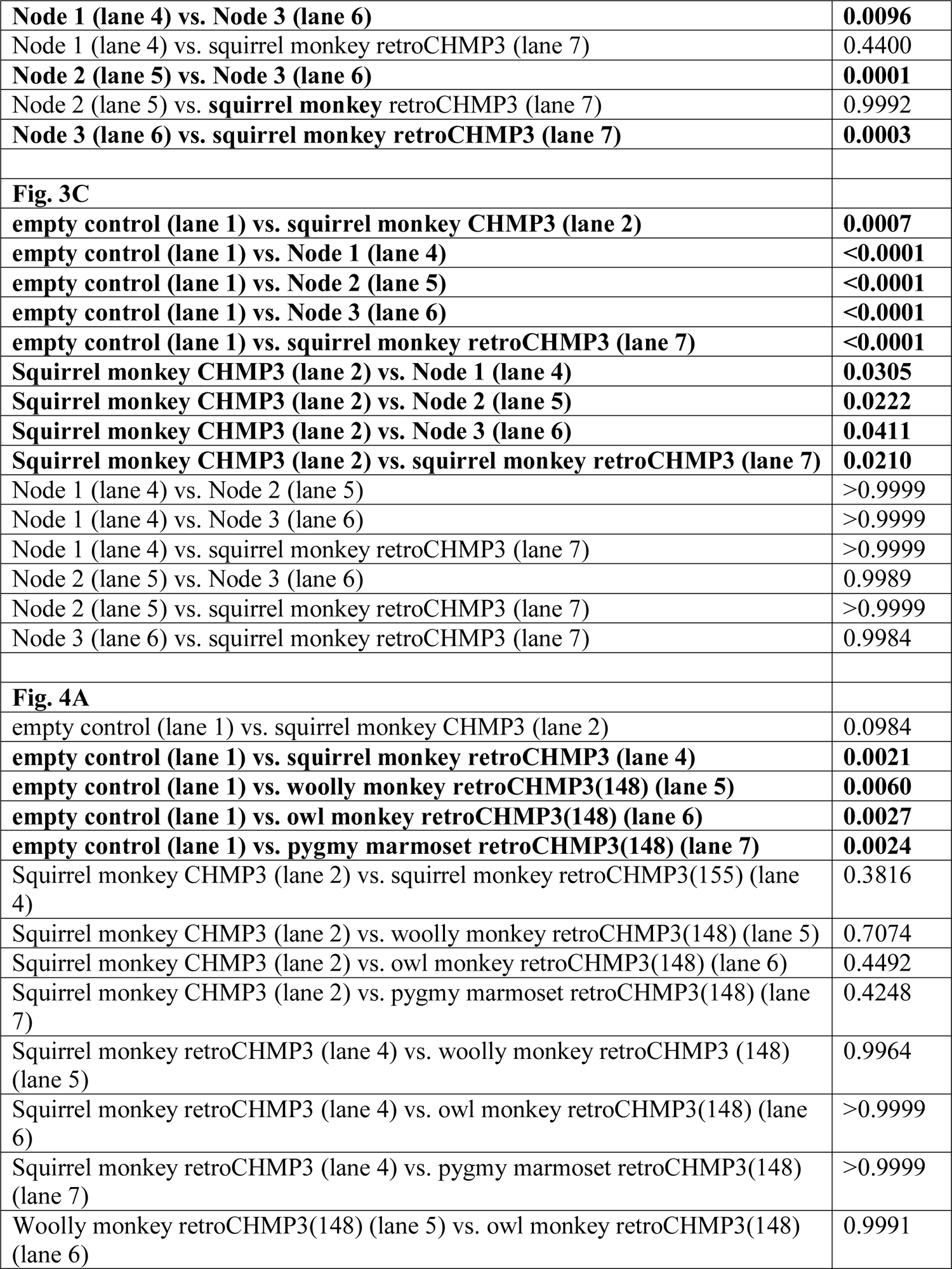
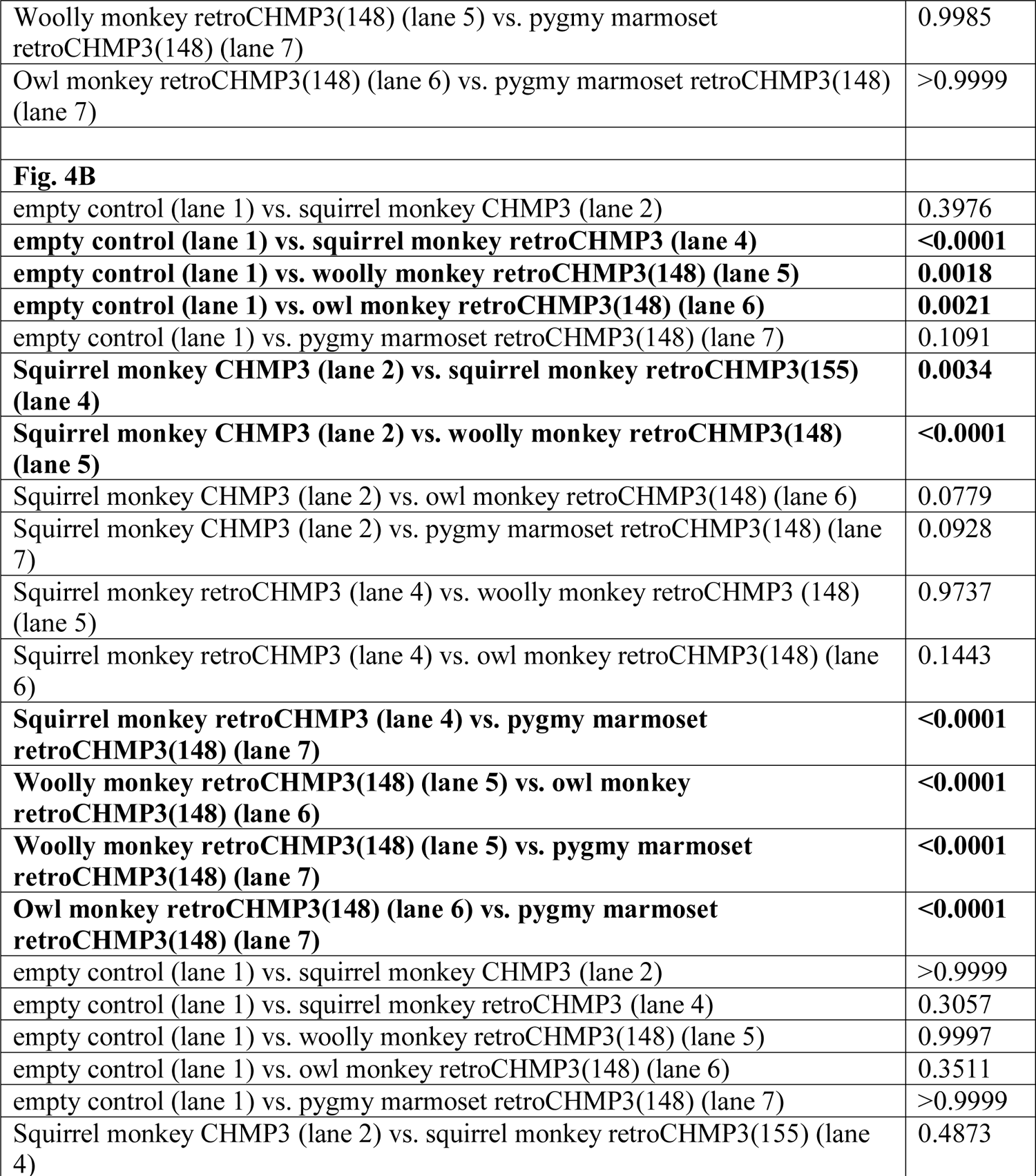
P-values for pairwise comparison of means for Fig. 2B, 2C, 3B, 3C, 4A and 4B. P-values were calculated using an ordinary one-way ANOVA followed by a Tukey’s multiple comparisons test.

**Table S7:**
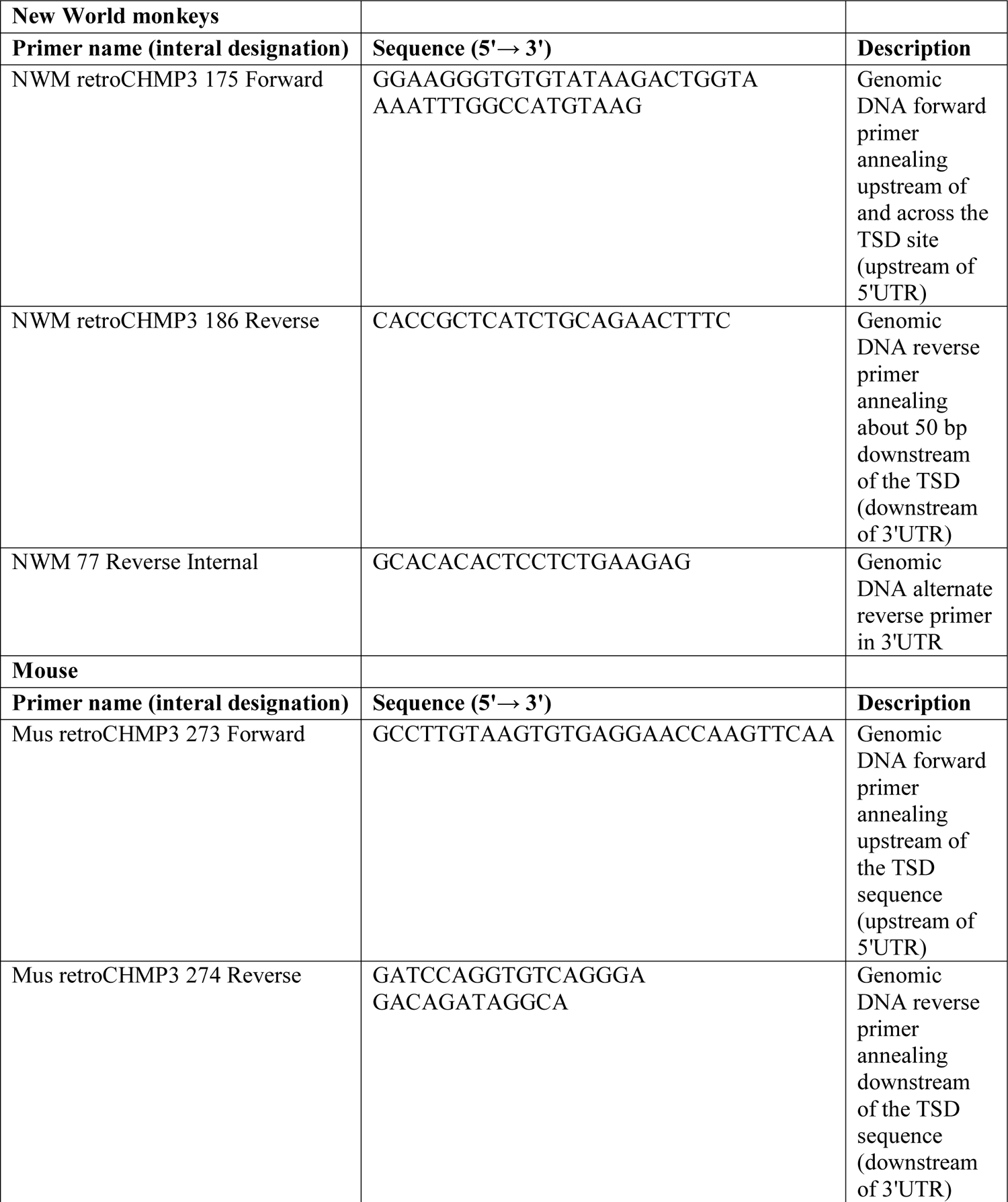
Primer sequences used in this study

## Materials and Methods

### Cells

HEK293T cells (CRL-3216, Human embryonic kidney endothelial cells, sex: female), GSML cells (CRL-2699, squirrel monkey B lymphoblast cell line, sex: male), and SML cells (CRL-2311, squirrel monkey lymphocytes cell line, clone 4D8, sex: male), were obtained from ATCC. MT-4 cells (Human T cells, NIH-ARP 120, sex: male) were obtained from the NIH AIDS Reagent program. Primary fibroblast cell lines for the following New World monkey species were obtained from Coriell Cell Repositories: *Saimiri sciureus* (common squirrel monkey, AG05311, sex: female), *Pithecia pithecia* (white-faced or Guianan saki, PR00239, sex: male), *Aloutatta sara* (Bolivian red howler monkey, PR00708, sex: male), *Saguinus fuscicollis* (Spix’s saddle-back tamarin, AG05313, sex: female), *Callicebus moloch* (dusky titi, PR00742, sex: female), *Callithrix geoffroyi* (white-fronted marmoset, PR00789, sex: female), *Lagothrix lagotricha* (common woolly monkey, PR00525, sex: female), *Aotus nancyma* (NancyMa’s owl monkey, PR00627, sex: male).

HEK293T were maintained in Dulbecco’s modified Eagle’s medium (DMEM, Gibco, Thermo Fisher Scientific) containing 10% FBS at 37°C and 5% CO_2_. Primary fibroblast cell lines were maintained in Dulbecco’s modified Eagle’s medium (DMEM, Gibco, Thermo Fisher Scientific) containing 10% FBS and penicillin/streptomycin at 37°C and 5% CO_2_. MT-4 and squirrel monkey B lymphoblast cells were maintained in RPMI-1640 medium (Gibco, Thermo Fisher Scientific) containing 10% FBS (Atlanta Biologicals) at 37°C and 5% CO_2_. Cells were tested for mycoplasma contamination every 3 months using the PCR Mycoplasma Detection Kit (abm) and were negative. Cells have not been authenticated.

### Plasmids

The expression constructs used in this study are listed in the Key Resources Table. All newly generated expression constructs have been submitted to the Addgene plasmid repository (https://www.addgene.org/). The HIV-1 expression construct NLENG1-IRES-GFP was a kind gift from David N. Levy (New York University) (Levy et al., 2004).

### Primate and mouse genetic sources

Sources for genomic DNA were as follows: Blood samples from *Cebus apella* (tufted capuchin, three individuals) were obtained from the NIH Animal Center (Dickerson MD, USA). Blood samples from *Ateles geoffroyi* (black-handed spider monkey, one individual) and *Ateles fusciceps* (brown-headed spider monkey, one individual) were obtained from Utah’s Hogle Zoo (Salt Lake City UT, USA). Blood samples from *Saimiri boliviensis boliviensis* (Bolivian squirrel monkey, 6 individuals), *Saimiri boliviensis peruviensis* (Peruvian squirrel monkey, 6 individuals) and *Saimiri sciureus sciureus* (Guyanese squirrel monkey, 6 individuals) were obtained from the Insitute Pasteur de la Guyane (Cayenne Cedex, French Guiana). Genomic DNA for the following species were obtained from Coriell Cell Repositories: *Saimiri sciureus* (common squirrel monkey, AG05311), *Pithecia pithecia* (white-faced or Guianan saki, PR00239), *Callicebus moloch* (dusky titi, AG06115), *Lagothrix lagotricha* (common woolly monkey, NG05356), *Saguinus labiatus* (red-chested mustached tamarin, AG05308), *Callithrix pygmaea* (pygmy marmoset, PR00644).

Sources for mouse genomic DNA were as follows: Blood samples from *Mus musculus* (house mouse, C57BL/6) were obtained from the University of Utah animal facility (Salt Lake City, UT, USA). Genomic DNA from *Mus spretus* and *Mus castaneus* were obtained from Guy Lenk (University of Michigan) and genomic DNA from *Mus dunni, Mus spretus, Mus musculus, Mus spicilegus, Mus minutoides* and *Mus pahari* were obtained from Christine Kozak (NIAID).

### Identification, sequencing and cloning of CHMP3 retrocopies

Retrocopies of CHMP3 were identified using NCBI BLAST (https://blast.ncbi.nlm.nih.gov/Blast.cgi). Specifically, the full-length CHMP3 cDNA (5’ UTR, coding sequence and 3’UTR) encoded by *Homo sapiens* (ENST00000263856.9, https://www.ncbi.nlm.nih.gov/nuccore/NM_016079.4) or *Mus musculus* (ENST00000263856.9, https://www.ncbi.nlm.nih.gov/nuccore/NM_025783.4) were used to query mammalian genomes on the UCSC Genome browser or whole genome shotgun contigs via NCBI BLAST to retrieve retrogenes present in the corresponding genomes. Candidate retrocopies were identified as CHMP3 sequences lacking introns, which is a key structural characteristic of processed, retrotransposed pseudogenes. In all cases, we have identified the 5’ and 3’ target site duplications flanking each retrocopy.

Retrogene insertions in New World monkeys and Mus genomes were validated by PCR amplification of genomic DNA using primers positioned in flanking sequences upstream and downstream of the target-site duplication. Primer sequences used are listed in Table S7. DNA was extracted from blood or cell lines using the Quick-DNA Miniprep kit (Zymo Research), following the manufacturer’s protocol. PCR products were resolved on 2% agarose gels followed by extraction and purification of candidate DNA amplicons using Zymoclean Gel DNA Recovery Kits (Zymo Research). Purified PCR products were cloned using the TOPO TA cloning kit (ThermoFisher Scientific), following the manufacturer’s instructions. Following transformation and plating, bacterial colonies were selected and grown in overnight cultures for plasmid purification using the Zyppy Plasmid Miniprep kit (Zymo Research). Sanger sequencing of clones was performed using M13F and M13R primers. RetroCHMP3 sequences are reported in Table S4 and S5.

### ORF decay modeling

To simulate the decay of retroCHMP3 under neutral selection, we used ‘mutator’ and ‘orf_scanner’ scripts (Young et al., 2018). The ORF of the ancestral new world monkey retroCHMP3 (666 bp) and the ORF of *Mus musculus* CHMP3 (GenBank accession NC_000072.6, 672 bp) were used as the starting ORFs. The following mutation rates (per site per generation, assuming 3 generations per year) were used for mouse (Uchimura et al., 2015; Yang et al., 2020): substitution = 5.4e^-9^, insertion/deletion: 1.55e^-10^. The following substitution rates were used for new world monkeys, and were based on mutation rates for owl monkey (*Aotus nancymaae*) and a generation time of one year (Thomas et al., 2018): substitution = 8.1e^-9^, insertion/deletion = 1.7e^-10^. Standard deviations for each mutation rate were calculated based on raw published mutation rate data (Thomas et al., 2018; Uchimura et al., 2015). ORF status was simulated (10,000 replicates) every 50,000 years for 50,000,000 years. The number of ORF that were still open and greater than 100 amino acids were counted every 50,000 years and divided into bins of full-length (224 or 222 aa), 140–180 aa or 180–220 aa.

### Ancestral reconstructions

Nucleotide sequences for New World monkey parent CHMP3 and retroCHMP3 were obtained by sequencing as described above. The parent CHMP3 sequences were aligned using the Multiple Sequence Alignment by Log-Expectation (MUSCLE) program (Edgar, 2004). The highest quality consensus sequence resulting from the alignment was generated in Geneious (https://www.geneious.com) and used as an approximation of the ancestral retroCHMP3 retrocopy (consensus parent). The consensus parent and retroCHMP3 sequences were aligned using MUSCLE. The alignment was manually edited to remove all insertions and deletions. The resulting edited alignment was used to infer the maximum likelihood phylogeny with RAxML (Stamatakis, 2014) using the GTR+GAMMA model of rate heterogeneity. The consensus parent sequence was specified as the outgroup. Statistical support for each node was evaluated by bootstrap analysis. The ancestral sequence reconstruction was performed using FastML (Ashkenazy et al., 2012). The edited multiple sequence alignment and ML phylogeny were used as inputs and the Jones-Taylor-Thornton model of amino acid substitution was used.

### HIV-1 budding assays

0.8 × 10^6^ HEK293T cells were seeded in 6-well plates 18-24 h before transfection. Cells were co-transfected using 10 μL Lipofectamine 2000 (Thermo Fisher Scientific) according to manufacturer’s instructions with 1 μg HIV-1 NLENG1-IRES-GFP expression vector and increasing amounts of expression vectors for full-length CHMP3 or retroCHMP3 to generate an expression gradient. Plasmid DNA amounts were as follows: For Fig. 2B, 1.5 μg pEF1α-squirrel-monkey-retroCHMP3-HA; 50 ng, 100 ng and 250 ng pEF1α-ancestral-retroCHMP3-HA; 250 ng, 500 ng and 1 μg pEF1α-owl-monkey-retroCHMP3-HA; for Fig. 3C, 100 ng pEF1α-squirrel-monkey-full-length-CHMP3-HA, 500 ng pEF1α-squirrel-monkey-CHMP3(155)-HA, 1.5 μg pEF1α-squirrel-monkey-retroCHMP3-HA, 2 μg pEF1α -NWM-Node 1, 2 μg pEF1α -NWM-Node 2, 2 μg pEF1α -NWM-Node 3; for Fig. 4A, 100 ng pEF1α-squirrel-monkey-retroCHMP3-HA, 1.5 μ -woolly-monkey-retroCHMP3(148)-HA, 1.5 μ pEF1α ygmy-marmoset-retroCHMP3(148)-HA, 1.5 μ pEF1α l-monkey-retroCHMP3(148)-HA. As necessary, plasmid amounts were adjusted with empty pCMV(WT) vector. The medium was replaced with 2 ml DMEM 4-6 h later. Cells were harvested 24 h post-transfection for western blot analysis, and supernatants were harvested for titer measurements and western blot analysis as described below.

For western blot analyses, virions from 1 ml supernatant were pelleted by centrifugation L 20% sucrose cushion (90 min, 15,000×*g*, 4°C) and denatured by adding 50 μL1x Laemmli SDS-PAGE loading buffer and boiling for 5 min. Cells were washed in 1 ml PBS, μL Triton lysis buffer (50 mM Tris pH 7.4, 150 mM NaCl, 1% Triton X-100) supplemented with mammalian protease inhibitor (Sigma-Aldrich). 150 μL 2× Laemmli SDS-PAGE loading buffer supplemented with 10% 2-mercaptoethanol (Sigma-Aldrich) were added, and samples were boiled for 10 min. Proteins were separated by SDS-PAGE, transferred onto PVDF membranes and probed with antibodies. Primary antibodies were as follows: anti-HIV CA (Covance, UT415), anti-HIV MA (Covance, UT556), anti-HA (Sigma-Aldrich, H6908), anti-GAPDH (EMD Millipore, MAB374). Antibodies with UT numbers were raised against recombinant proteins purified in the Sundquist laboratory. Bands were visualized by probing the membrane with fluorescently labeled secondary antibodies (Li-Cor Biosciences) and scanning with an Odyssey Imager (Li-Cor Biosciences). HIV-1 titers were assayed in MT-4 cells by quantifying GFP expression in target cells.

Briefly, MT-4 cells (100,000 cells/well, 96-well plate) were infected with three different dilutions of virus-containing culture media in duplicate. After 16 h, dextran sulfate was added to μg/ml. Cells were harvested after 48 h, washed in PBS three times and fixed with 2% paraformaldehyde for 20 min at room temperature. Percentages of GFP-positive cells were determined by flow cytometry on a FACSCanto (BD Biosciences).

### Toxicity assays

For cytotoxicity assays, 1.5 × 10^4^ HEK293T cells were seeded in 96-well plates. After 24 h, cells were transfected with Lipofectamine 2000 (Thermo Fisher Scientific) according to manufacturer’s instructions. Each well received varying amounts of HA-tagged CHMP3 construct, shown to result in equal expression across all constructs by Western blot. After 48 h, cellular toxicity was analyzed using the CellTiter-Glo Luminescent cell viability assay (Promega) following manufacturer’s instructions. Luminescence was detected using a Biotek Synergy microplate reader.

To test protein expression levels, 5 × 10^5^ HEK293T cells were seeded and transfected in parallel in 6-well plates. After 48 h, cells were harvested, pelleted and lysed in Pierce RIPA Buffer (Pierce) containing Halt Protease Inhibitor (Thermo Fisher Scientific). Samples were mixed with equal amounts 2x Laemmli Sample Buffer containing 10% 2-mercaptoethanol and heated at 95°C for 30 min. Samples were run on Mini-Protean TGX™ Precast Protein Gels (Biorad) and transferred to Immobilon-P PDVF Membrane (Millipore). Membranes were blocked in PBS + 0.1% Tween 20 + 5% powdered milk for 1 h and then incubated with anti-HA (Sigma-Aldrich, H6908) or Anti-Actin monoclonal antibody made in mouse (ThermoFisher Scientific AM4302) for 2 h at room temperature. After three washes in PBS + 0.1% Tween 20, membranes were incubated in Anti-Rabbit IgG HRP conjugate (Millipore #AP132) or Anti-Mouse IgG/IgM HRP conjugate (Millipore #AP130P) for 2 h, washed three times in PBS + 0.1% Tween 20 and developed with Advansta WesternBright ECL HRP substrate. Blots were imaged using a LI-COR blot scanner.

## Statistical analysis

Statistical significance for budding assays and toxicity assays were determined by using a one-way ANOVA followed by Tukey’s multiple comparison test, using GraphPad Prism 8 software.

## Notes

### Competing Interest Statement

The authors have declared no competing interest.

